# Ganglioside enriched phospholipid vesicles induce cooperative Aβ oligomerization and membrane disruption

**DOI:** 10.1101/2022.04.14.488413

**Authors:** Jhinuk Saha, Priyankar Bose, Shailendra Dhakal, Preetam Ghosh, Vijayaraghavan Rangachari

## Abstract

A major hallmark of Alzheimer disease (AD) is the accumulation of extracellular aggregates of amyloid-β (Aβ). Structural polymorphism observed among Aβ fibrils in AD brains seem to correlate with the clinical sub-types suggesting a link between fibril polymorphism and pathology. Since fibrils emerge from a templated growth of low-molecular weight oligomers, understanding the factors affecting oligomer generation is important. The membrane lipids are key factors that influence early stages of Aβ aggregation and oligomer generation, and cause membrane disruption. We have previously demonstrated that conformationally discrete Aβ oligomers can be generated by modulating the charge, composition, chain length of lipids and surfactants. Here, we extend our studies into liposomal models by investigating Aβ oligomerization on large unilamellar vesicles (LUVs) of total brain extracts (TBE), reconstituted lipid rafts (LRs) or 1,2-dimyristoyl-sn-glycero-3-phosphocholine (DMPC). Specifically, we varied the vesicle composition by varying the amount of GM1 gangliosides added as a constituent. We found that liposomes enriched in GM1 induce the formation of toxic, low-molecular weight oligomers that are isolable in a lipid-complexed form. Importantly, the data indicate that oligomer formation and membrane disruption are highly cooperative processes. Numerical simulations on the experimental data confirm cooperativity and reveal that GM1-enriched liposomes form twice as many numbers of pores as those without GM1. Overall, this study uncovers mechanisms of cooperativity between oligomerization and membrane disruption under controlled lipid compositional bias, and refocuses the significance of the early stages of Aβ aggregation in polymorphism, propagation, and toxicity in AD.

**TOC figure:** 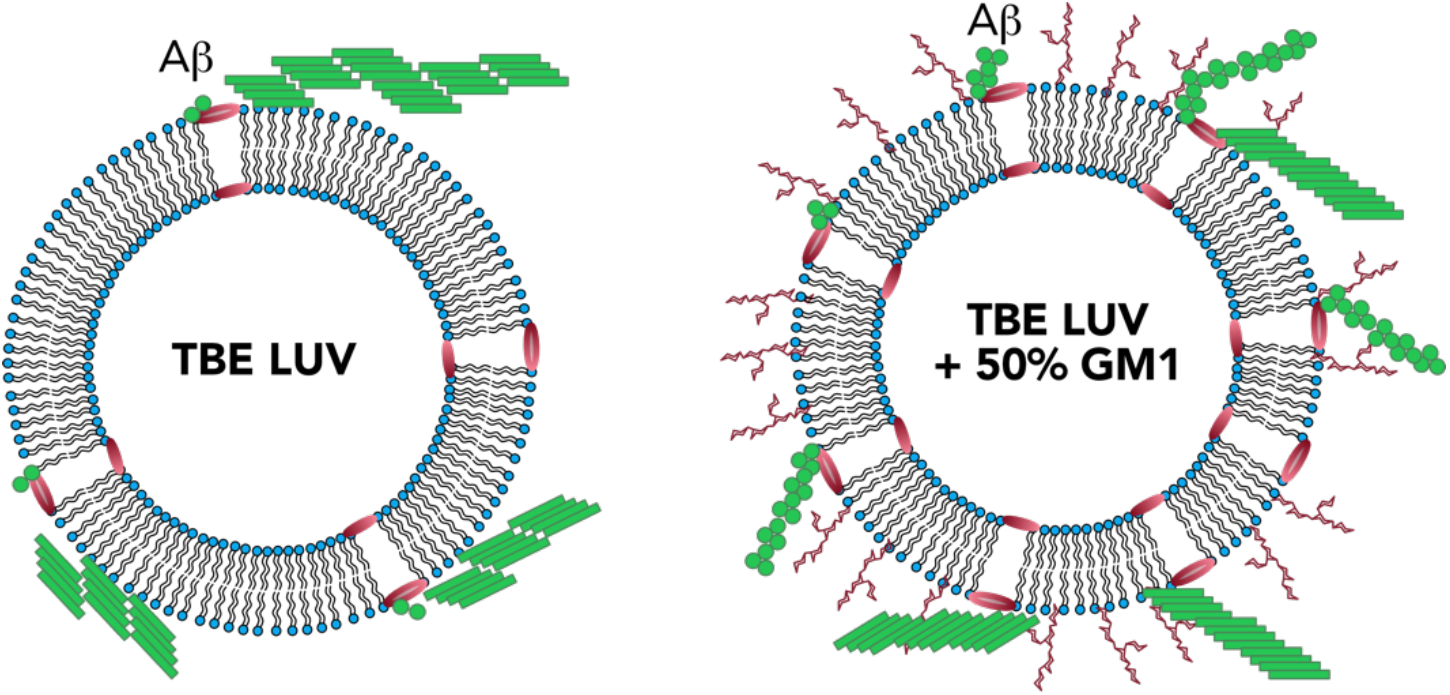

## INTRODUCTION

Alzheimer disease (AD) is a neurodegenerative disorder associated with deposition of extracellular plaques composed of amyloid-β (Aβ) aggregates in the brain. Aβ peptide is generated by the sequential cleavage of transmembrane amyloid precursor protein (APP) by β and γ secretases and is subsequently released into the extracellular space^1–3^. Monomeric Aβ is intrinsically disordered and undergoes near spontaneous aggregation towards high molecular weight insoluble fibrils involving a sigmoidal growth kinetics ^4–6^. The low molecular weight soluble oligomers generated during aggregation are known to be the primary toxic species in early stages of AD pathology that impair hippocampal synaptic plasticity and cause blockage of long-term hippocampal potentiation (LTP) ^7–10^. A few mechanisms by which the oligomers impart toxicity are, membrane disruption via pore formation, release of reactive oxygen species (ROS), astrocytosis and microglial activation^11–14^. It has been long hypothesized that being closely associated with Aβ, membrane lipids and surfactants are likely to interact and generate conformationally diverse low molecular weight oligomers^13,15-18^. Lipids play an important role in the early stages of Aβ aggregation that dictates oligomer generation ^19–21^. We demonstrated that micelle forming lipids including fatty acids, lysophospholipids and gangliosides can induce distinct conformational oligomers that have discrete cellular and pathological functions ^22^. Many of these oligomers are toxic to neuroblastoma cells^19^ and induce cerebral amyloid angiopathy (CAA) in transgenic CRND8 mice^18^.

Extensive investigations in the past have revealed that the kinetics and structural dynamics of Aβ aggregation is influenced by membrane components and constitution. Liposomes containing anionic phospholipids, sphingomyelins and sterols have been reported to cause rapid amyloid formation^23–26^. Furthermore, the aggregation rates of Aβ have are modulated differently depending on the surface charges on small unilamellar vesicles (SUVs) containing negatively charged phosphoglycerol (PG), neutral phosphocholine (PC) or on large unilamellar vesicles (LUVs) containing a mixture of PC/PS or PC/PG lipids^27,28^. In addition, other membrane components such as cholesterol and gangliosides have also been known to influence membrane Aβ interaction^29,30^. Accelerated membrane disruption by Aβ has been observed in ganglioside containing model membrane systems^14^. Aβ has been observed to preferentially bind to regions containing GM1 in raft-like lipid vesicles enriched with GM1 and cholesterol and augment aggregation ^25,31-36^, and morphologically distinct Aβ fibril polymorphs have been known to form in the presence of GM1 containing model vesicles^37^. Furthermore, cell membrane and its components also facilitate membrane disruption and pore formation by Aβ aggregation^14,38,39^. However, since the formation of low-molecular weight oligomers are influenced the most by lipids, it remains unclear whether oligomerization and membrane disruption are discrete events that are temporally decoupled from one another or the two have synergistic relationship. To address this question, here we enriched GM1 ganglioside in varying amounts on LUVs and SUVs of 1,2-dimyristoyl-sn-glycero-3-phosphocholine (DMPC), reconstituted lipid rafts (LR), and total brain extract lipid (TBE) to see the dynamics of Aβ42 (referred from here on as Aβ) oligomerization and membrane disruption. We observed that high percentage of GM1 ganglioside doping generates distinct low molecular weight oligomers of Aβ that can be isolated and characterized. More interestingly, oligomerization and membrane disruption are cooperative. Numerical simulations uncover that GM1 doping forms trimeric oligomers that form pores, which further assists aggregation of oligomers toward high molecular weight species. Addition of preformed aggregates to the vesicles however, forms pores in a more abrupt manner. These results provide new mechanistic insights on the possible role of gangliosides on the membrane surface towards synergistic Aβ oligomerization and toxicity.

## RESULTS

### TBE and LR LUVs enriched with GM1 ganglioside promote the formation of Aβ oligomers

First, to obtain insights into the effect of GM1 ganglioside enriched vesicles on the temporal dynamics of Aβ aggregation, freshly purified, seed-free Aβ monomers (25 μM) buffered in 20 mM Tris (pH 8.0) containing 50 mM NaCl and 50 mM thioflavin-T (ThT) were incubated with 0.3 mg/mL preprepared LUVs of DMPC, LR or TBE individually at 37 °C. The three liposomal systems were chosen such that to capture diverse set of membrane compositions. The liposomes were made by systematically increasing the amount of GM1 gangliosides added (% by weight) from 0 to 50%. The aggregation kinetics was monitored by ThT fluorescence on a 96-well plate reader. The control Aβ in the absence of liposome (◁ in Figure 1a, b, and c, respectively) followed a typical sigmoidal pattern with a lag time of ~5 hours. Surprisingly, incubation of Aβ with LUVs of DMPC without GM1 showed similar or slightly decreased lag time to that of Aβ in the absence of vesicles (■ Figure 1a). Incubation of Aβ with LR or TBE LUVs without GM1 gangliosides showed decreased lag times of 2-3 hours (■ Figures 1b and c). However, LUVs enriched with increasing amounts of GM1 ganglioside showed significant decrease in lag times and increase in fluorescence intensity within two hours of incubation (○, ▲, ▽ & ♦ for 10, 25, 33, and 50% GM1 doping respectively; Figures 1a, b and c). With micellar systems, we have previously reported the generation of discrete Aβ oligomer^18^. Therefore, to investigate whether similar oligomer generation is facilitated by GM1-enriched LUVs, the incubated reactions were monitored by immunoblotting in parallel. The samples from the reactions in Figure 1a, b and c were electrophoresed under partial denaturing conditions after 3, 5 and 9 hours of incubation, and visualized via immunoblotting using the monoclonal antibody Ab5. Aβ incubated with unenriched LUVs showed monomeric, dimeric, and trimeric bands after 3 h (lane ‘0’; Figure 1d, e, and f). After 5 h and 9 h the dimeric and trimeric bands disappeared with a concomitant appearance of high molecular weight bands that failed to enter the gel that are possibly fibrils

**Figure 1:**
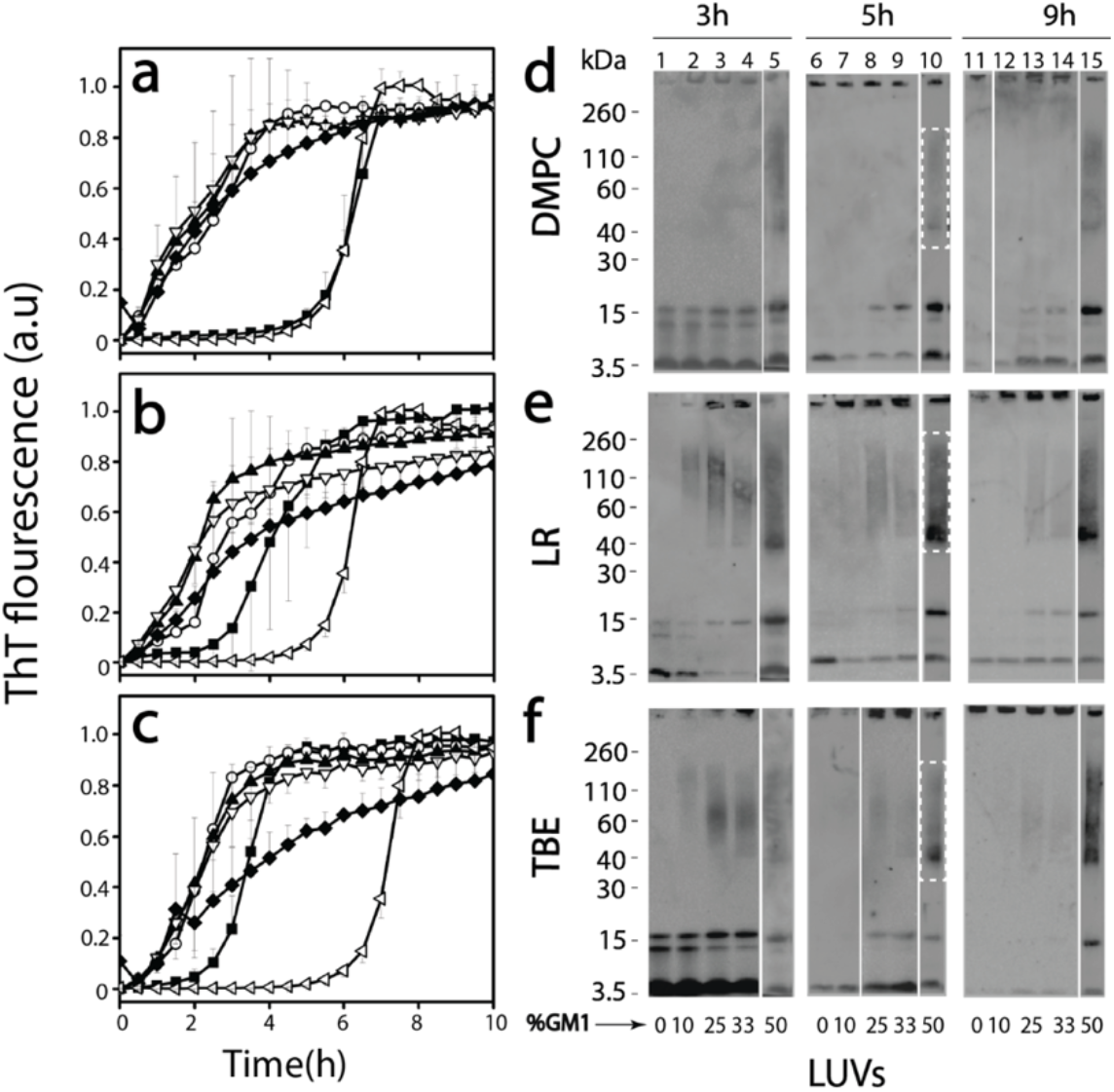
(a, b, and c) Normalized ThT fluorescence kinetics of buffered 25 μM Aβ without (◁ control) or with DMPC (a), LR (b) and TBE (c) LUVs each of them enriched with 10 (○), 25 (▲), 33 (▽), and 50 (♦) % GM1 ganglioside (by wt.) or without (■) GM1 in presence of 50 mM NaCl in 20mM tris buffer pH 8.00. (d, e, and f) Partially denaturing SDS-PAGE immunoblots of 25 μM Aβ in presence of DMPC, LR and TBE LUVs respectively, enriched without or with 10, 25, 33, and 50% GM1. Gels were run at intervals of 3, 5 and 9 hours, respectively.

(5 and 9 h, 0 %; Figure 1 d, e, and f). Similarly, incubation of Aβ with increasing amounts of GM1 also showed dimer and trimer bands along with monomers in case DMPC LUVs (Figure 1d: 10, 25 and 33%) upon 3h of incubation. Transition from dimer and trimer to higher molecular weight fibrils has been observed to decrease with the increase in GM1 percentage of the LUVs. Furthermore, faint oligomeric bands ranging from 40 to160 kDa emerged after 5h of incubation (50%; Figure 1d) which were stable till 9h of incubation (Figure 1d; *lane* 15). Immunoblots of Aβ incubated with increasing GM1 enriched LUVs in LR and TBE showed dimer and trimer band for LUVs with lower GM1 content (Figure 1e and f). The intensity for these oligomeric bands were greater for 50% GM1 containing LR and TBE LUVs as compared to 25 or 33%. Also, these oligomers were present up to 9h of incubation (Figure 1e and f). In all samples, bands near 4.6 and >260 kDa were also observed, which indicate the presence of monomers and high molecular weight fibrils, respectively (Figure 1d, e, and f). These results suggest that increase in GM1 ganglioside content in vesicles influence the generation of Aβ oligomers.

### Secondary structure transitions during aggregation reveal potential conformational intermediates specific to GM1 enrichment

To investigate conformational changes of Aβ during aggregation, far UV CD spectroscopy was used. Samples containing LUVs with no, or enriched with 50% GM1 gangliosides from Figure 1 were analyzed. To see whether there are differences in the early oligomer formation among different LUVs due to change in their surface characteristics, we monitored the reaction for the initial five hours. In all reactions as expected, Aβ showed conformational conversion from a random coil to β-sheet upon aggregation (Figure 2), consistent with the ThT fluorescence and immunoblot results in Figure 1. Aβ incubated with DMPC LUVs enriched with 50% GM1 showed an immediate conversion from random coil (λ^min^ = 200 nm) to β-sheet (λ^min^ = 218 nm; dark blue region in the contour plot) (Figure 2a), while those with no GM1 showed a slow conversion from a persistent random coil structure to β-sheet (Figure 2b), also consistent with ThT aggregation kinetics. LUVs of LRs enriched with 50% GM1 cause a more rapid transition of random coil to β-sheet than the unenriched ones (Figure 2c and d). Aβ incubated with LUVs of TBE however, show a gradual transition from a random coil to α-helical within the first 1.5 hours followed by the transition to a β-sheet signal (Figure 2e and f). The α-helical intermediate was more apparent in TBE LUVs enriched with 50% GM1 (Figure 2e). Among the GM1 enriched vesicles, DMPC showed the slowest transition from random coil to β-sheet and TBE was the only one in which an α-helical intermediate was observed.

**Figure 2.**
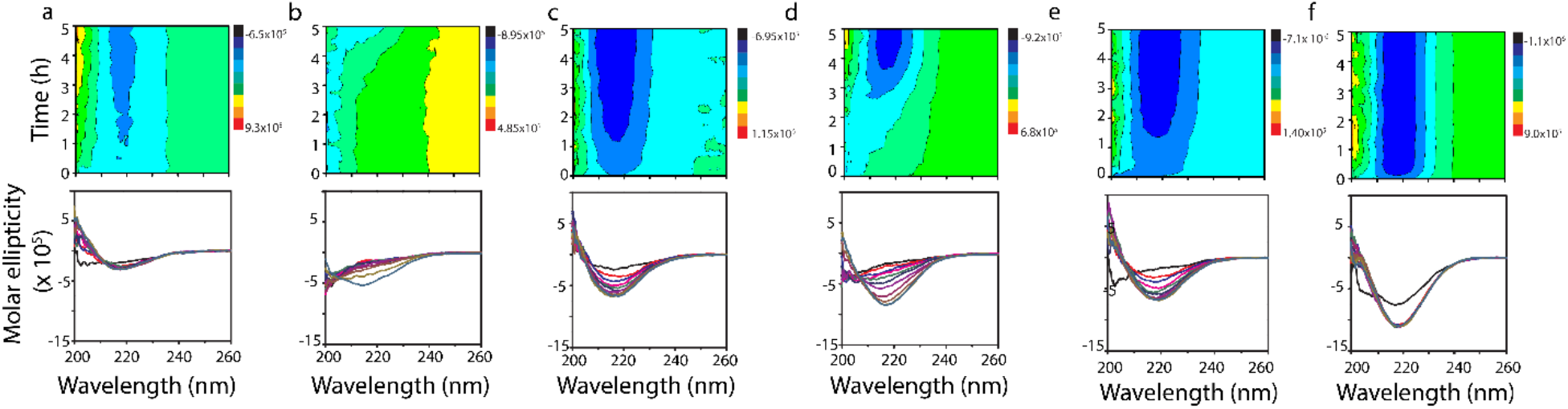
Far-UV CD contour and time course plots for buffered (20 mM tris buffer pH 8.0, 50 mM NaCl) 25 μM monomeric Aβ incubated with LUVs of 50 and 0% GM1 enriched DMPC (a and b, respectively), 50 and 0% GM1 enriched LR (c and d, respectively), and 50 and 0% GM1 enriched TBE (e and f, respectively) collected for up to five hours at 37 °C in quiescent conditions.

### Aβ oligomers isolated from GM1 enriched vesicles show distinctive biophysical characteristics

To see if oligomers generated in the presence are isolable, freshly purified Aβ monomers (25 μM) were incubated with 50% GM1 LUVs (DMPC, LR and TBE respectively) at 37 °C quiescent conditions. To isolate the oligomeric species from the reactions containing monomeric or fibrillar species, samples after 5 hours were centrifuged at 18000xg for 20 minutes. The supernatant was then subjected to fractionation by size exclusion chromatography (SEC) using a Superdex-75 column. Fractionation of all three LUVs showed two peaks near the void volume with a small first peak at fraction 15 and a larger one between fractions 16 and 18 (solid line; Figure 3c). In addition, a third peak at an included volume at fraction 24 was observed (solid line; Figure 3c).

**Figure 3.**
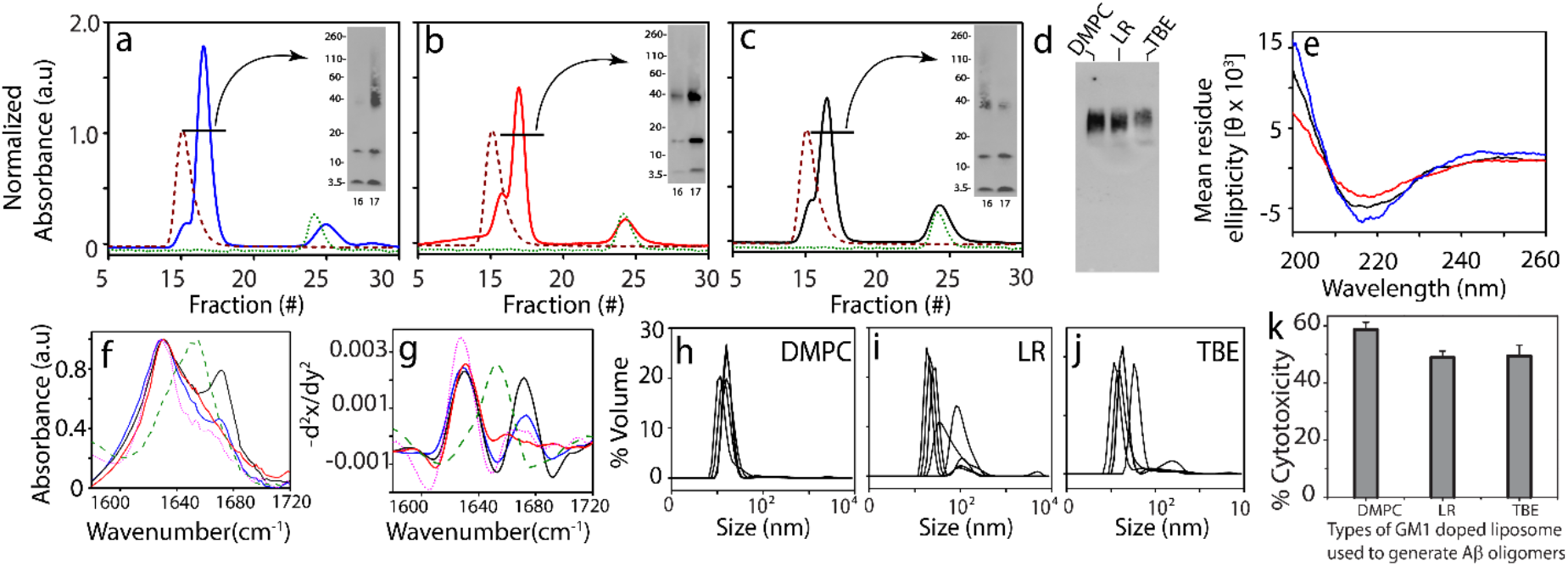
(a-c) SEC chromatogram for isolation of Aβ oligomers generated in presence of 50% GM1 enriched DMPC 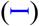, LR 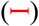and TBE LUVs 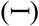 respectively, LUV control at 0.3 mg/mL 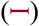 and control Aβ 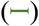 at 5 h, inset-SDS PAGE immunoblots of SEC isolated oligomer fraction 16-17 (d) Native PAGE immunoblot for SEC isolated Aβ oligomers generated in the presence of 50% GM1 enriched DMPC, LR and TBE LUVs respectively, (e) CD spectra of fraction 17 of SEC isolated Aβ oligomers generated in the presence of 50% GM1 enriched DMPC 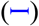, LR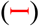 and TBE LUVs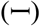 respectively (f) FTIR spectra for SEC isolated Aβ oligomers generated in the presence of 50% GM1 enriched TBE 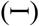, DMPC 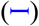 and LR 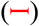 LUVs, homotypic Aβ fibril 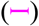 and BSA control 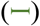 respectively (g) Negative of double derivative of the FTIR spectra (fig3. (f)) (h-j) DLS for fraction 17 of SEC isolated Aβ oligomers generated in the presence of 50% GM1 enriched DMPC, LR and TBE LUVs respectively, (k) XTT assay performed on SHY5Y neuroblastoma cells upon incubation with isolated Aβ oligomers from 50% GM1 enriched DMPC, LR and TBE LUVs respectively expressed in terms of % of dead cells. n=3 independent cell cultures on isolated oligomers, statistically significant at p< 0.05 based on one-way ANOVA analysis.

The first fraction at 15 corresponded to free vesicles (purple dashed line; Figure 3c) while the fraction at 24 corresponded to monomeric Aβ (green dotted line; Figure 3c). After 5 h of incubation, the aliquots of the fractions 16 and 17 were subjected to electrophoresis under partial denaturing conditions (with 1% SDS and without sample boiling) and visualized by immunoblotting (Figure 3a-c; insets). In all samples, monomeric bands near 4.6 kDa, multiple oligomeric bands near 15 kDa, and 38-110 kDa as well as high molecular weight fibrils bands which could not enter the gel were visible in the immunoblots. Fractions from DMPC showed more disperse oligomers between 38 and 110 kDa (Figure 3a) while those from LR and TBE showed more compact band pattern centered around 38 kDa corresponding to ~ 8mer. To see whether the low molecular weight oligomeric bands observed were due to dissociation of the oligomers due to SDS treatment during electrophoresis, the isolated oligomeric samples were run under nondenaturing conditions (no SDS, no sample boiling) in PAGE gel followed by immunoblotting. By doing so, homogenous oligomeric bands without the presence of any other band were observed suggesting that the monomeric and other low oligomer bands were probably due to dissociation of oligomers by SDS treatment (Figure 3d). The secondary structure of isolated oligomers was investigated by far-UV CD and FTIR spectroscopy. All oligomers were found to have β-sheet structure evident from minima at 217 nm in far UV-CD with the exception of those derived from LR which showed a small extent of helical structure (shoulder at 222 nm) (Figure 3e). Similarly, the amide I band of the FTIR signature was investigated to gain more insights about the type of β-sheet (parallel or anti-parallel) within the oligomers generated with GM1 enriched LUVs. The absorbance maxima for all three oligomer samples showed a band near 1630 cm^-1^ without a 1690 cm^-1^ band indicative of a parallel β-sheet structure ^40^ (Figure 3f). However, only oligomers generated with TBE and DMPC LUVs enriched with 50% GM1showed a second band near 1671cm^−1^ (Figure 3 f and g), which is indicative of turn conformation^41,42^. It can be inferred that TBE catalyzed oligomers have some structural differences compared to those from LR or DMPC generated LUVs, which parallels the observation of conformational transitions with TBE LUVs (Figure 2). Size of isolated oligomers analyzed by DLS revealed that these oligomers are 18-20 nm in diameter (Figure 3 h-j). However, presence of polydispersity in these oligomers suggest the possible co-elution of some amounts of LUVs with the oligomers. Indeed, we found that only about 0.05 mg/mL (~ 17%) of the starting amount of lipids remain associated with the isolated oligomer (Figure S3). Furthermore, Aβ oligomers were tested for their toxicities on SHY5Y neuroblastoma cells by XTT assay^22^. All three oligomers were toxic with 50% cell viability. DMPC generated oligomers had a slightly higher cytotoxicity compared to LR and TBE (Figure 3k). Overall, these data suggest that LUVs with different surface properties and charge could lead to the generation of structurally distinctive neurotoxic oligomers as observed in micellar systems^22^.

### GM1 enriched vesicles induce cooperative Aβ oligomerization and membrane pore formation

Aβ incubated with the LUVs of TBE enriched with 50% GM1 showed the presence of possible conformationally different oligomer intermediate (Figure 2 and 3). To further investigate whether these oligomers also induce membrane pore formation, dye leak assay was performed using 6-carboxyflourescein (6-FAM) encapsulated within TBE vesicles. Freshly purified Aβ monomers (10 μM in 20 mM Tris pH 8.0) were incubated with 6-FAM loaded TBE LUVs and fluorescence was monitored in a 96 well plate for 12 h at 37 °C (see Methods). Aβ monomers when incubated with TBE LUVs without GM1 showed no discernable membrane disruption (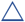; Figure 4a). However, when incubated with 50% GM1-enriched TBE LUVs, Aβ monomers showed increased FITC fluorescence at ~ 2 h of incubation that continued to steadily increase up to 20% during the next 9 h (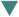; Figure 4a) suggesting steady disruption of the vesicles. In contrast, preformed fibrils isolated from Aβ incubations with TBE LUVs without or with 50% GM1 showed exponential increases in pore formation (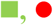; Figure 4a). A similar patten was observed when the same fibrils were sonicated (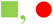; Figure 4b).

**Figure 4:**
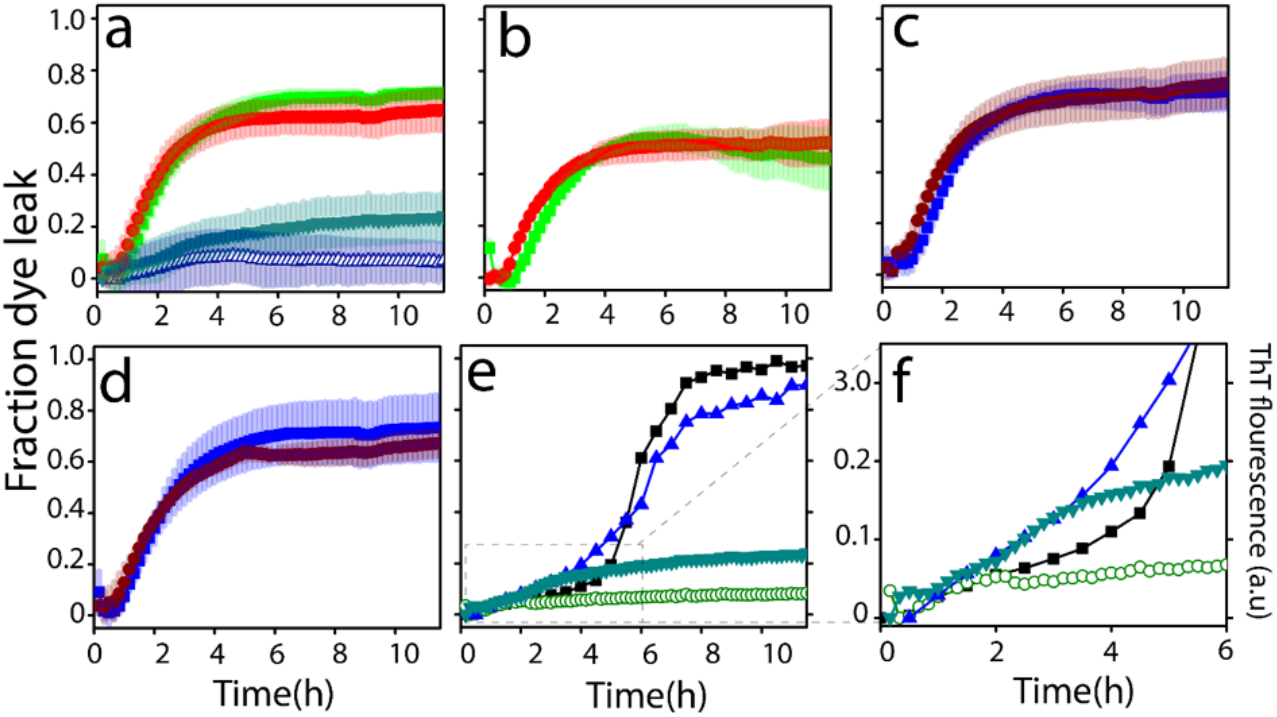
Vesicle dye leak analysis monitored by 6-carboxyflourescein (6-FAM) dye on; (a) TBE LUVs incubated with 10 μM Aβ monomers 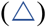 or 2 μM isolated Aβ fibrils generated from the same liposomes 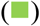; 50% GM1-enriched TBE LUVs incubated with 10μM Aβ monomers 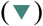; or 2 μM isolated Aβ fibrils generated from 50% GM1-enriched liposomes 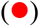; (b) TBE LUVs incubated with 2 μM sonicated Aβ fibrils generated in the presence TBE liposomes 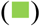 or 50% GM1-enriched TBE LUVs incubated with 2 μM sonicated Aβ fibrils generated in the presence of 50% GM1-enriched liposomes 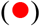; (c) TBE LUVs incubated with 2 μM isolated Aβ fibrils generated in the absence of liposomes 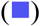 or 50% GM1-enriched LUVs incubated with 2 μM isolated Aβ fibrils generated in the absence of liposomes 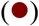; (d) samples in (c) but sonicated; (e) ThT fluorescence of 10μM Aβ monomers in presence of 50% GM1-enriched TBE LUVs 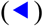 and 50% GM3-enriched TBE LUVs (■); 6-FAM dye leakage of 50% GM1-enriched TBE LUVs 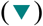 and 50% GM3-enriched TBE LUVs 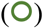 in presence of 10μM Aβ monomers; (f) zoomed-in image of figure 4(e) showing the initial 6 h of the reaction.

When fibrils generated from Aβ in the absence of liposomes were incubated on TBE LUVs without or with 50% GM1 showed exponential increase in pore formation either unsonicated (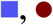; Figure 4c) or sonicated (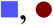; Figure 4d). Together, it is evident that high molecular weight fibrils are able to disrupt the membranes more efficiently than low molecular weight oligomers but there are several possible caveats as discussed further below. Nevertheless, it is evident that GM1 ganglioside enrichment promotes oligomers vis-à-vis membrane disruption as opposed to unenriched liposomes. To specifically see whether glycoform distributions on the gangliosides have an effect on these properties, as we had seen before with Aβ-glycopolymer interactions^43,44^, TBE liposomes were also enriched with GM3 gangliosides which have significant sugar distribution differences with GM1 (Figure S2). By doing so, none or minimal leakage of dye upon incubation of Aβ monomers with 50% GM3 enriched TBE LUVs (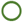; Figure 4e) was observed. When both the dye-leak data and ThT data with GM1- and GM3-enriched LUVs are compared, the specificity of interactions is evident; while GM3 enriched liposomes showed a sigmoidal pattern of aggregation without pore formation, reactions with GM1-enriched samples showed an aggregation and concomitant pore formation during the first 3 h (Figure 4e and f).

### Numerical simulations uncover insights into the cooperativity in oligomerization and membrane disruption

To obtain more details on the effects of GM1 on Aβ oligomerization and membrane disruption, ordinary differential equation (ODE) based numerical simulations were used. The basis of the models along with the reaction abstractions formulated are detailed in the Methods section. We used Scatter Search optimization algorithm to fit the experimental data as it has been earlier shown that metaheuristic algorithms like scatter performs better than other algorithms to fit the Aβ aggregation ^45^. Sum of squared errors (SSE) was used as a metric to evaluate the models, and COmplex PAthway SImulator (COPASI) ^46^ to solve the mathematical models. Briefly, oligomerization was considered up to the formation of 12mers, beyond which all aggregates were considered ‘fibrils’ for modeling simplicity. An additional reason was to identify the low molecular weight oligomeric species that are responsible for membrane disruption and not those that were formed late. Individual global fits of the ThT and the FITC dye-leak data of Aβ aggregation on TBE liposomal with varying gangliosides were performed. Specifically, the modeling was directed at understanding the temporal mechanisms and cooperativity by which Aβ aggregated and caused membrane disruption as a function of GM1 enrichment of liposomes.

To do so, two potential pathways of pore formation upon Aβ oligomerization on membrane surfaces were considered. Upon aggregation that generates a single pore, Aβ can then elongate/aggregate on the edge of the pore assisted by the exposed membrane components. This can either result in further enlargement of the pore or aggregates could initiate the second pore formation and so on. Since both mechanisms involve cooperativity, we arbitrarily chose the latter mechanism to model and due to lack of experimental evidence for either mechanism.

Model simulations based on the rate constants computed (Tables S1-S5). A global fit of the ThT aggregation and FITC dye leak data a good fit and an agreement with the experimental data (Figure 5a-c). The models showed that an increase in GM1 percentage results in more pores on the membrane surface. For example, it was found out that in the case of liposomes with 0% GM1, only two pores are formed whereas, for enrichment of the TBE liposomes with 50% GM1 showed that four different pores were formed. In other words, enrichment with 50% GM1 resulted in twice as many numbers of pores as those without GM1. Computation of various aggregate sizes formed temporally during aggregation suggested that dimeric Aβ was responsible for pore formation in the absence of GM1 while trimeric Aβ was responsible for 50% enriched GM1 liposomes. In our reaction system, the smallest and the largest oligomers were considered to be 2 and 6mers for the control in the absence of GM1 and 2 and 8 for GM1-enriched liposomes as the lower and upper bounds. The oligomer responsible for causing the initial pore was computed by sweeping the oligomer size to fit the FITC data; this gave the least SSE (Tables S1-S5) for dimer (for 0% GM1 control) and 3mer (for 50% GM1). Furthermore, cooperativity in pore formation and aggregation was also evident from the rate constants obtained. It can be observed that for 0% GM1, 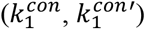 is less than 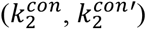 suggesting higher cooperativity especially for 50% GM1, which showed greater cooperativity in both creation of subsequent pores and in aggregation of the oligomers. This aspect of cooperativity separates the mechanism by which LUVs in the absence of GM1 form pores but do not promote robust pore-forming fibrils. However, the concentrations of the oligomers responsible for pore formation during aggregation were low in the order of ~ 0.5 μM at 2-3 h of incubation (Figure 5d-f), that explains the difference in the rates of pore formation between the preformed fibrils and oligomers generated in situ.

**Figure 5:**
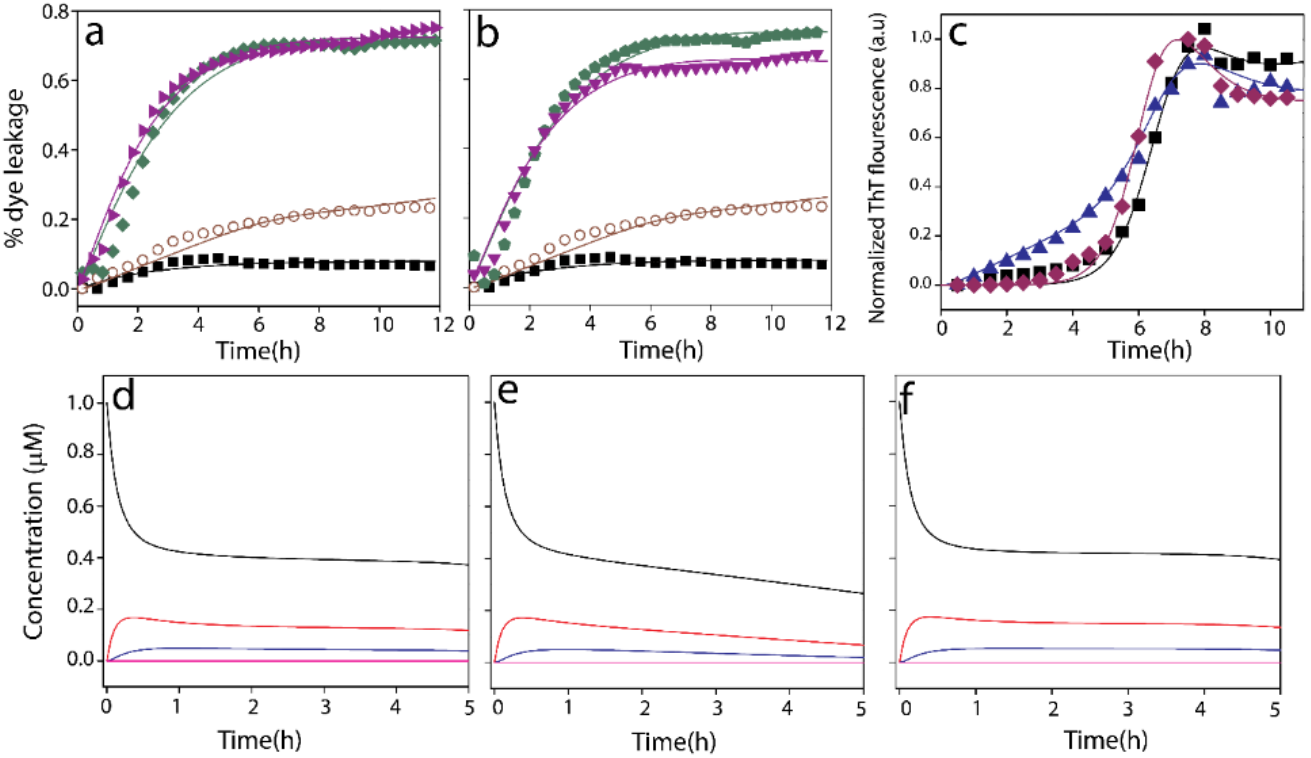
Computational fits of 6-carboxyflourescein dye leak assay of TBE LUVs (a) with 50% GM1 and 10 μM Aβ monomers 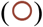, without GM1 and 10 μM Aβ monomers (■), with 50% GM1 and 2 μM Aβ fibrils 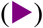, without GM1 and 2 μM Aβ fibril 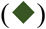 (b) with 50% GM1 and 10 μM Aβ monomers 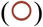, without GM1 and 10 μM Aβ monomers (■), with 50% GM1 and 2 μM sonicated Aβ fibrils 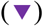, without GM1 and 2 μM sonicated Aβ fibril 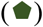 (c) Normalized ThT fluorescence kinetics of buffered 10 μM Aβ without (■; control) or with TBE LUVs each of them enriched with 50 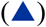 % GM1 ganglioside (by wt.) or without 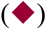 GM1 in presence of 50 mM NaCl in 10mM sodium phosphate buffer pH 8.00; Aβ monomers (A1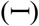), oligomers (A2 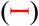 and A3 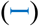) and fibrils (F 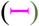) distribution plots for first 5 h from the start of reactions of Aβ monomers with TBE LUVs, (d) no GM1, (e) 50% GM1 or (f) without LUVs.

## DISCUSSION

Aggregate polymorphism is increasingly becoming known as a distinguishable feature among many AD patients^47–53^. It is speculated that such polymorphic fibrils are in part responsible for the observed phenotypes. Since fibrils are the end products of templated aggregate growth, we hypothesize that the conformational differences and selection among low molecular weight oligomers are keys in determining the dominant fibril polymorph. In this regard, we have previously shown that membrane lipids and surfactants modulate Aβ aggregation pathways to generate conformationally distinct oligomers capable of propagating their structure towards fibrils^18,54^. Specifically, we have observed that Aβ oligomers generated in the presence of lipid micelles are structurally distinct and cause distinct phenotype in APP transgenic mice^18^. Similarly, a plethora of studies point towards the effect of other membrane model systems like liposomes on aggregation of Aβ and membrane disruption ^55–60^. For example, properties of membranes such as surface charges^20,61^ curvature^62,63^, composition ^64^ etc., have been shown to have profound effects on aggregation. Similarly, Aβ aggregates are known to form pores and channels in the membrane that are attributable to their biophysical characteristics^14,38,65,66^. However, it remains unclear whether and how low molecular weight Aβ oligomers are generated upon its interaction with liposomal surfaces and whether such a generation is dependent on the membrane components. Furthermore, the coupling between oligomerization and membrane pore formation remains unclear.

The study presented here shows alteration of surface characteristics, especially to the degree of charge density by dilution with neutral GM1 gangliosides seem to decisively affect the oligomerization of Aβ (Figure 6). While LUVs without or very low amount of GM1 accelerates the aggregation of Aβ to form higher molecular weight fibrils in the first five hours of incubation, LUVs enriched with high concentration of GM1 causes oligomerization of Aβ on the LUV surface, kinetically trapping the Aβ oligomers. Among the three different types of LUVs used, i.e., DMPC, LR and TBE that have different compositions, were found to augment aggregation of Aβ but also showed oligomerization when enriched with 50% GM1. This implicates the significance of gangliosides in Aβ aggregation as previous studies have established^35,67-70^. Furthermore, the GM1 enriched TBE LUVs showed somewhat modified ThT aggregation kinetics that correlated with a partially helical conformational state at an early aggregation stage. More importantly, these temporal changes also coincided with membrane disruption brought upon only by high GM1-enriched samples. This phenomenon may be due to altered lipid-packaging or dilution of anionic charge density or both due to GM1 enrichment. It is noteworthy that the pore formation was not abrupt but rather was slow and progressive in nature but only showed ~ 20% at the end of 11 hours of incubation (Figure 4). By contrast, fibrils and sonicated fibrils of Aβ generated in the presence and absence of liposome showed rapid pore formation. Furthermore, addition of fibrils generated from GM1-enriched liposomes too showed rapid pore formation. Two possible explanations can be rendered for these observations; *a*) the oligomers formed during aggregation is low in concentration as computed (~ 0.5 μM; Figure 5e) to effect rapid change in pore formation kinetics, or *b*) not oligomers but high molecular weight fibrils effect membrane disruption more effectively. A third explanation could be that the mechanisms of pore formation could be either numerous small pores or a few large pores for monomers aggregating on the surface or when pre-formed aggregates are added, respectively. Yet another key observation is that the oligomerization and membrane disruption is also selective to the nature of sugar distributions on gangliosides. While GM1 ganglioside promotes membrane pore formation, GM3 does seem to have such an effect, nor does it promote oligomers. Collectively, the data indicate that for early stages of oligomer formation, membrane selectivity is important, to generate conformationally distinct and toxic species, however at later stages when the higher molecular weight species are already formed, membrane is ruptured in a different mechanism than while Aβ oligomerization. Furthermore, oligomerization and pore formation seem to be cooperative and coupled to one another. As mentioned earlier, our lab and others have reported the formation of structurally distinct Aβ aggregates with equally distinct biophysical properties in presence of GM1 gangliosides. Recently, Matsuzaki and his group reported the formation of amyloid tape fibrils with mixed parallel and antiparallel β-sheet structure in presence of GM1 in membrane model systems^68^. Therefore, it can be concluded that membrane lipid composition as well as GM1 content plays a role in generating distinct oligomers. This inference is further supported by our CD time course data, where secondary structure of the intermediate species along the pathway of oligomerization are different for liposomes enriched with GM1 gangliosides. In this report, we further these findings to uncover that GM1 ganglioside enrichment of membrane vesicles not only promotes oligomerization but also induces membrane disruption in a cooperative manner. This suggests that aggregation and modulation of membrane dynamics can be coupled to each other, one that is facilitated as a function of specific membrane compositions. Such cooperative mechanisms may lead to the generation of conformational oligomers, which we believe are key templates for the formation of polymorphic fibrils observed in patient brains.

**Figure 6.**
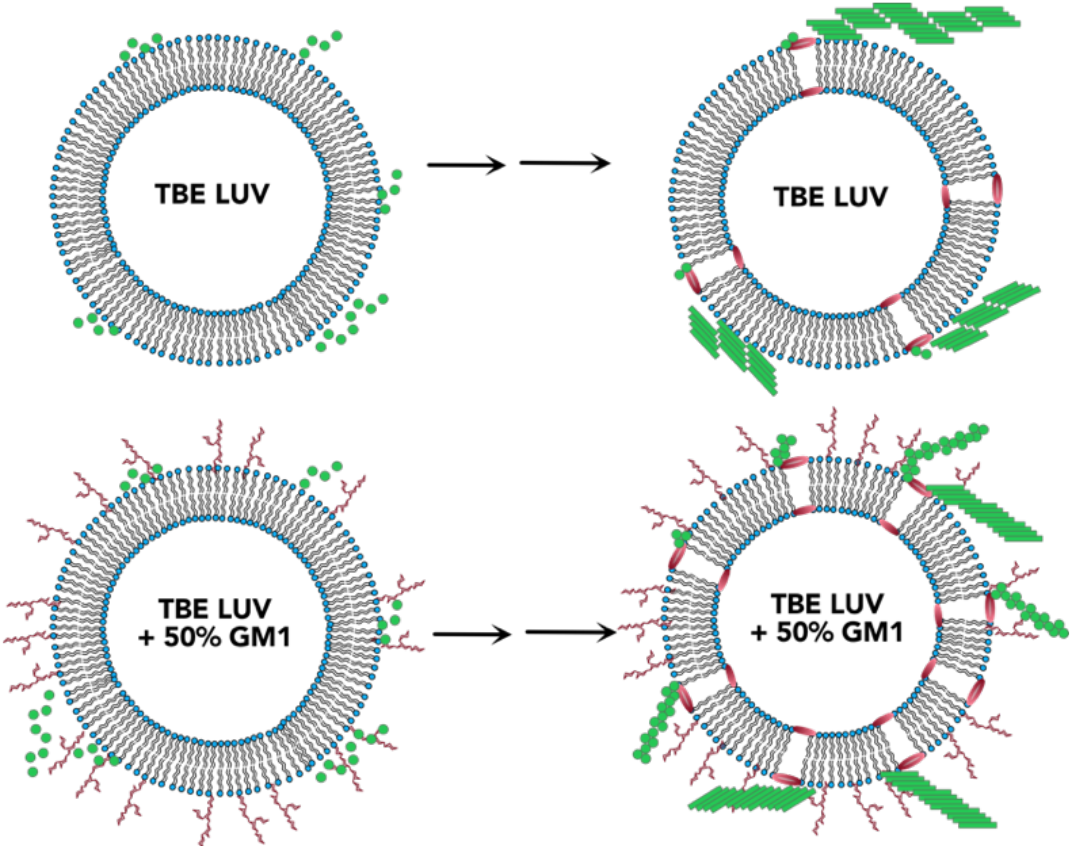
Schematic of conclusions drawn from this study showing the effect of GM1 enrichment in liposomes.

## EXPEREMENTIAL PROCEDURES

### Materials

Size exclusion chromatography (SEC) column (Superdex-75 HR 10/30) was purchased from GE Life Sciences (Marlborough, MA). DMPC, 1-palmitoyl-2-oleoyl-glycero-3-phosphocholine (POPC), 1-palmitoyl-2-oleoyl-glycero-3-phosphoethanolamine (POPE), sphingomyelin, cholesterol and total brain lipid extract (TBE) were purchased from Avanti Polar Lipids, Inc. (Alabaster, AL); Tris base, Tris hydrochloride, and SDS were purchased from Sigma-Aldrich (St. Louis, MO) or Thermo Fisher Scientific, Inc. (Waltham, MA). Other chemicals, reagents, and consumables were purchased from either VWR, Inc. (Radnor, PA) or Thermo Fisher Scientific, Inc. (Waltham, MA). The monoclonal antibody Ab5 was obtained from Dr. Levites at the University of Florida (Gainesville, FL). Liposome extrusion system was purchased from Avanti Polar Lipids, Inc. (Alabaster, AL). The plasmid, pET-Sac Aβ(M1-42) was obtained from ADDGENE.

### Recombinant Aβ expression and purification

Recombinant Aβ (Aβ(M1-42)) was recombinantly expressed in BL21(DE3) PlysS Star *E. coli* cells. Cells were grown in LB broth and induced for 16h and subsequently harvested and lysed by sonication to obtain inclusion bodies. Inclusion bodies were resuspended in 6M urea and filtered with 0.2μm hydrophilic PVDF filter. Filtrate was directly subjected to HPLC chromatography using a Zorbax C8 column pre-heated at 80°C. Purified Aβ was lyophilized and stored at −80 °C for further use^71^. To obtain monomers, HPLC purified Aβ (0.5 −1 mg) was resuspended in 490 μL of nanopure water and allowed to stand for 30 mins. NaOH was then added to the mixture to a final concentration of 10 mM and was allowed to stand for 10 minutes at room temperature. The mixture was then loaded onto a Superdex-75 HR 10/30 SEC column pre equilibrated with 20mM Tris pH-8.00 and attached either to an AKTA FPLC system (GE Healthcare, Buckinghamshire) or a BioLogic DuoFlow™ system (BioRad) fractionating at a flow rate of 0.5 mL/min at 25 °C. Monomers were eluted between fractions 24 and 28. Molar concentration of each monomer fraction was determined by UV absorbance collected using a Cary 50 UV-Vis spectrometer (Agilent Technologies, Inc.; Santa Clara, CA) and subsequently applying Beer-Lambert’s law ε = 1450 cm^−1^ M^−1^ at 276 nm). Purity and integrity of the peptide was confirmed using matrix-^72^assisted laser desorption/ionization (MALDI) time-of-flight mass spectrometry. Purified monomers were stored in 4°C and used for experiment within the same day of purification.

### Liposome preparation

LUVs were prepared as done previously ^14,67,73-75^.DMPC, POPC/POPE/ sphingomyelin/cholesterol in 33/33/10/20 percent by weight (for LR), and TBE liposomes were constructed from a 1:1 chloroform: methanol solution of lipids stocks. The solution was gently dried under nitrogen flow and then placed in a vacufuge with desiccant overnight to further evaporate any residual solvent. The dried lipid film was then either rehydrated with a either buffer solution (10 mM phosphate or 20 mM tris buffer, pH 8.0) or buffer solution containing doping agent 10 to 50 percent (by weight) of GM1or GM3 for DMPC or LR & TBE respectively, to yield a final concentration of 1 mg/mL. The hydrated lipids were shaken vigorously for 1-1.5 h at 37 °C and subjected to 15 freeze thaw cycles with liquid nitrogen and hot water at ~50 °C. The resulting solution was extruded 25 times through a 200 nm (for LUVs) polycarbonate nucleopore membrane filter (Whatman) with a mini extruder to obtain unilamellar vesicles. The size of the vesicles was confirmed with DLS collected using a Zetasizer Nano S instrument (Malvern, Inc., Worcestershire, UK) as described below.

### Thioflavin-Tkinetics

Aβ monomers (25 μM or 10 μM) was incubated with 0.3 mg/ml DMPC/ TBE / raft-like reconstituted (LR) LUVs/SUVs in either 20mM tris or 10 mM sodium phosphate buffer (pH 8.00) in presence of 50mM NaCl and 50 μM ThT. Kinetics were read in corning black 96-well plates in a Biotek Synergy well plate reader at 37 °C monitored for every 30 min with shaking for 10s before every read. The fluorescence data was processed and normalized from 0 - 1 using Origin8.0 as done earlier^76^.

### Isolation of oligomers

Aβ oligomers were generated by incubating freshly purified Aβ monomer (25μM) with the specified LUVs/ SUVs in the conditions listed below. 0.3 mg/mL DMPC LUVs; 0.3 mg/mL lipid raft LUVs; 0.3mg/ml TBE LUVs. Additionally, 50 mM NaCl were added to all reactions prior to incubation at 37 °C in quiescent conditions for 5h. The samples were then pelleted by centrifugation at 18000g for 20 min and the soluble supernatant was subjected to SEC as described above. Fractions of 500 μL were collected, and Aβ oligomers were found to be in the 16-17^th^ fraction. The molar concentration after isolation was determined by UV-vis spectroscopy, as described above. Samples were either stored at 4 °C and used for experimentation within 72 h or lyophilized and kept at −80 °C for extended storage prior to experimentation. The size of the oligomers was confirmed with DLS.

### Electrophoresis and immunoblotting

Samples were run in denaturing SDS page gel by diluting samples in 1X Laemmli loading buffer, without boiling, onto either 4-12% NuPAGE or 4-20% Bis-Tris BioRad TGX gels. For molecular weight determination pre-stained molecular-weight markers (Novex Sharp Protein Standard, Life Technologies) were run in parallel with samples on the gel. Proteins were transferred on to 0.2 μm immunoblot membrane (biorad) using a thermo scientific transfer cassette for 15min. Subsequently, the immunoblot with protein was boiled for 1 min in a microwave oven in 1X PBS, followed by blocking for 1.5 h at 25 °C in 1X PBS containing 5% nonfat dry milk with 1% Tween 20. Blots were then probed overnight at 4 °C with a 1:6000 dilution of Ab5 monoclonal antibody, which detects amino acids 1—16 of Aβ. Following primary incubation, blots were probed with a 1:6000 dilution of anti-mouse, horseradish peroxidase-conjugated secondary antibody for 1.5 h at 25 °C before being imaged using a Super Signal™ West Pico Chemiluminescent Substrate kit (Thermo Fisher Scientific).

### Dye leak assay

Lipid stocks, DMPC, POPE/POPC/sphingomyelin / cholesterol (in proportions described above for LR) and TBE stored in 1:1 chloroform:methanol was dried under liquid nitrogen and vacuufuged overnight as described previously ^14,74,75,77^and rehydrated with 15mM 6-carboxyflourscein (6 - FITC) in 10 mM sodium phosphate buffer pH 8.00. The rehydrated lipid-dye mixture was subjected to 15 freeze thaw cycles and subsequent extrusion with 200nm polycarbonate nucleopore membrane filter (Whatman) with on a mini extruder to obtain dye filled LUVs. The excess dye in the solution was separated from dye filled LUVs using 7 kDa desalting columns pre-equilibrated with 10 mM sodium phosphate buffer pH 8.00 centrifuged at 500xg for 30 sec. The size of the LUVs was confirmed using DLS as mentioned below. The leakage of dye was confirmed by comparing the fluorescence intensity (λ_EX_:490nm; λ_Em_:595nm) of intact dye-encapsulated liposomes and two-threefold increased intensity upon complete rupture of liposome upon addition of 0.2% TritonX-100 ^75^. The percent dye leak is calculated by the difference between the dye leak intensity of LUVs with the protein and blank divided by the difference between the dye leak intensity of LUVs with Triton X-100 and blank LUVs as done previously^14^.

### Dynamic light scattering (DLS) analysis

DLS was obtained with Zetasizer Nano S instrument (Malvern, Inc., Worcestershire, UK) by running a total of 15 runs for 10 sec each for every sample after an equilibration for 30 sec. The %volume function was used to calculate the diameter of the LUVs.

### Fourier transform infrared spectroscopy (FTIR)

FTIR was obtained with an Agilent FTIR instrument (Cary-630) with dial-path accessory. 45-50 μg of lyophilized protein samples (Aβ isolated oligomers/monomers) were resuspended in 5 μL D_2_O and samples were scanned from 1500-1800 cm^−1^ at a resolution of 4 cm^−1^. A total accumulation of 1024 spectral scans were obtained per sample and data were processed by subtracting the blank D_2_O spectra and baseline correction using OriginLab8.

### Circular dichroism (CD) spectroscopy

CD spectra of the oligomers/monomers were obtained using a Jasco (Easton, MD) J-815 spectropolarimeter. An average of 6-16 spectral scans in the far-UV region (260-190 nm) at a rate of 50 nm/min (8 s response time, 1 nm bandwidth, 0.1 nm data pitch). Savitzky-Golay algorithm with a convolution width of 15 was used to smoothen the spectra in the Jasco spectrum analysis program.

### Cell viability XTTassay

<u>Cell viability</u> was measured using 2,3-bis(2-methoxy-4-nitro-5-sulfophenyl)-5-[(phenylamino)carbonyl]-2H-tetrazolium hydroxide (XTT) assay kit (Biotium) using our previously established protocol^76^. Briefly, experiments were carried out in human neuroblastoma SH-SY5Y cell lines (ATCC) grown in DMEM and Ham’s F12K (1:1) medium containing 10% FBS and 1% Penicillin/streptomycin. Cells were maintained at incubator conditions set to a temperature of 37 °C and 5.5% CO2. Cells were seeded at a density of 30,000 per well in a clear bottom 96 well plate 24 hours prior to oligomers incubation. Oligomers were incubated at 2.5 μM concentration for 24 hours prior to performing XTT assay. All experiments were done in triplicates, statistical analysis and data processing was carried out using Origin 8.0.

### Model simulations

We have used a system of Ordinary Differential Equations (ODEs) to simulate and fit the experimental data as shown previously^54,78,79^. Parameter estimation is solved as an optimization problem in ODE systems by minimizing the objective function that calculates the deviation between simulated and experimental data. Optimization methods can be gradient-independent or gradient-based; the former method is theoretically less susceptible to the stochastic noise than the latter method. Hence, in this case of stochastic optimization, we have used gradient-free metaheuristics ^45,46^ as our algorithm for parameter estimation. The following modelling abstraction was used in this study.

#### a) Control (on-pathway) reactions

First, the ThT aggregation data for control Aβ (10 μM) in the absence of liposomes were mapped to concentration of the on-pathway fibrils. The forward and backward nucleation and the forward and backward fibrillation rate constants were calculated, and SSE was recorded. For modeling simplicity, aggregates beyond 12mers were considered to be fibrils and the rate constants were modeled for on-pathway fibril formation (Table S1), which is the basis for modeling other reactions. The following reactions were considered:

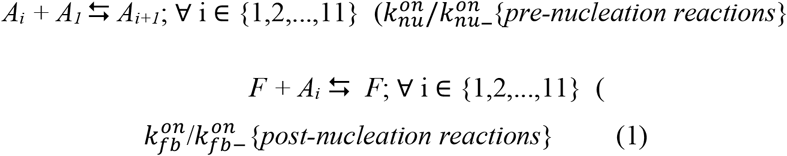

To reduce the number of species considered in the on-pathway reactions, we abstracted all post-nucleation species (*A*_12_ onwards) as on-pathway fibrils denoted by *F*. Here, the forward and backward rate constants (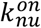 and 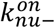, respectively) for all pre-nucleation reactions were considered the same to reduce the number of parameters and based on our prior work^18,78^. Similarly, the forward and backward rate constants (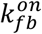 and 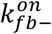, respectively) for all post-nucleation reactions were also considered the same. The intensity of the ThT data was mapped to the sum of the concentrations of the on-pathway fibrils as follows:

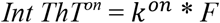

where, *k^on^* is a scaling constant used for fitting ThT intensity (*ThT^on^*) to the fibril concentration.

#### b) Reactions involving liposomes

Firstly, the forward and the backward pre-nucleation and post-nucleation rate constants computed from the controls were used for the on-pathway species for oligomerization reactions (shown below), which facilitated the reduction of the number of estimable parameters in this phase. To model the oligomerization reactions, both the ThT aggregation kinetics data and the FITC dye-leak data were considered. In our models, the initial concentrations of the liposomes were varied as their molar mass could not be precisely calculated. A sequential, multiple pore formation model was considered as opposed to one expanding pore although both are possible; since we do not have enough evidence to discount one over the other, we chose the former arbitrarily. Two possible scenarios were considered based on experimental evidence: pores formed by (*i*) a pre-nucleation oligomer (*A_j_*) and/or (*ii*) on-pathway fibrils (i.e., a post-nucleated oligomer denoted as *F*). These were modeled for the first pore, *BA_i_*, by;

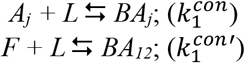

while for subsequent holes (*CA_i_, DA_i_, EA_i_*,), this is modeled by reactions of type:

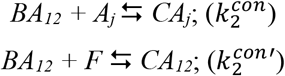

Here, *A_j_* denotes the minimum pre-nucleation oligomer that can form a pore and the value of the *i*mer was identified through our parameter fitting mechanism. Moreover, the values of the rate constant combination (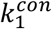 and 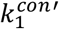) can suggest which mechanism is more likely for the first pore formation (i.e., through pre-nucleation oligomer or post-nucleation fibrils); similarly, the rate constant combination (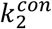 and 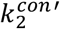) suggests which mechanism is more likely for the second pore formation and so on. The cooperativity between pore formation and aggregation was captured by considering further oligomerization reactions assisted by the edge of the pore up to 24mers denoted by reactions of type:

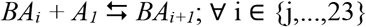

We additionally consider a bulk oligomerization in the presence of fibrils by reactions of type:

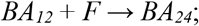

The cooperativity amongst pores is captured by (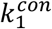 and 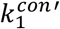) etc. For example, note that the second hole (*CA_i_*) is formed only after the first pore is formed. This is ensured by reactions of type:

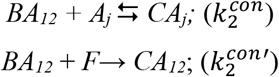

where, the presence of *BA_i_*, is necessary for the formation of *C_i_*. Additionally, if the rate constant pair (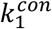 and 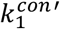)) is less than (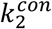 and 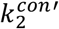)), this will suggest higher cooperativity in hole formation; in other words, the second hole formation (controlled by 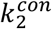 and 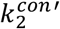) is faster than the formation of the first hole. Finally, to map the concentration values to the ThT and FITC signals, we considered all the species weighted by the oligomer size for FITC signal (denoted by summations ranging from 1,…,24) while only weighted values of post-nucleated oligomers were considered in the ThT signal (denoted by summations ranging from 12,…,24).

The reactions considered for the oligomerization phase are as follows:

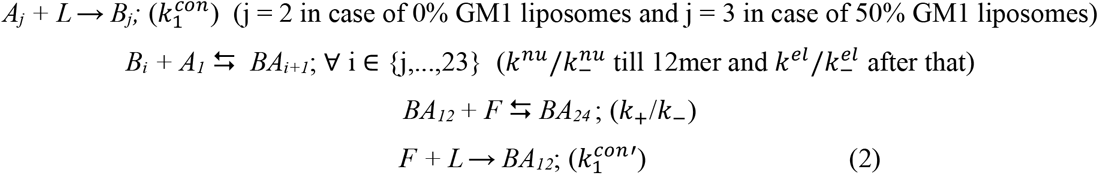

where *BA_i_*, denotes the first hole with an oligomer of size *i*-mers, and *L* is the liposome.

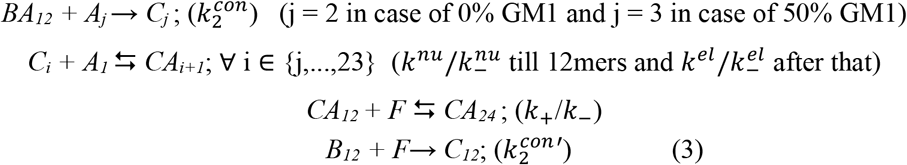

Fitting of data for 0% GM1 enriched liposomes were done only with the above reactions. Here, *C_i_* denotes the second hole with an *i*-mer. Similarly, for the following reactions, *D_i_* denotes the third hole, E*_i_* denotes the fourth hole and so on.

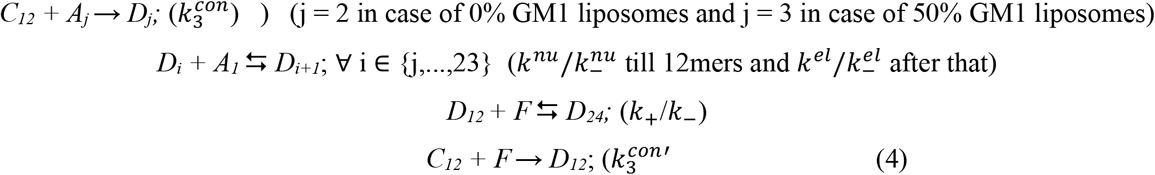

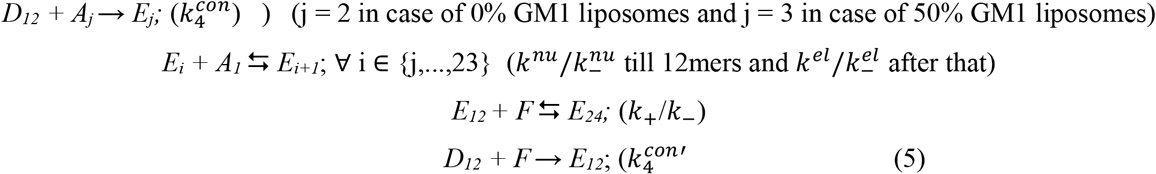

Fitting of data for 0% GM1 enriched liposomes were done only with the above reactions. i.e. four pores. Then ThT data were mapped to the sum of the concentrations of the on-pathway fibrils and all the off-pathway oligomers beyond 12mers as follows:

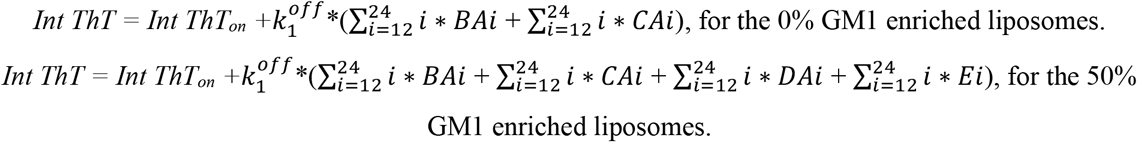

The FITC dye leak data were mapped to the concentration of the off-pathway oligomers as follows:

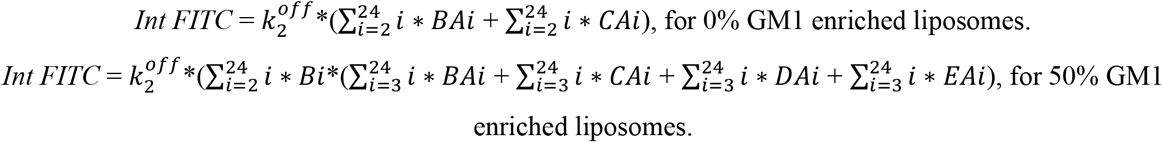

The model is extendable to any number of holes and the curve fitting with the experimental data infers the optimal number of holes to be considered. For example, for 0% GM1, we first experimented with four holes (*B_i_, C_i_, D_i_, E_i_*), which was then systematically reduced to one hole (*B_i_*); in this case two holes gave the best global fit with ThT and FITC dye leak data.

## Supporting information

none

## ACKNOWLEDGEMENTS

The authors would like to thank the following agencies for their financial support: National Institute of Aging (1R56AG062292-01) and the National Science Foundation (NSF CBET 1802793) to VR. The authors also thank the National Center for Research Resources (5P20RR01647-11) and the National Institute of General Medical Sciences (8 P20 GM103476-11) from the National Institutes of Health for funding through INBRE for the use of their core facilities.

## SUPPLEMENTARY INFORMATION

**Figure S1:**
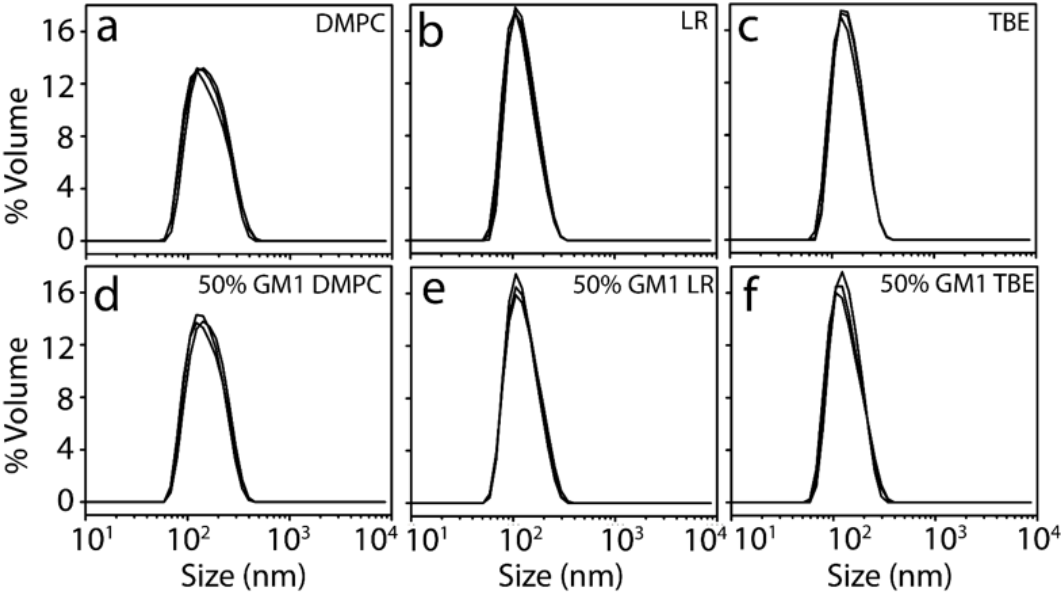
DLS of (0.2 mg/mL) DMPC, LR and TBE LUVs with (a,b, and c) or without (d,e, and f) GM1 extruded with 200 nm pore-sized polycarbonate membrane.

**Figure S2:**
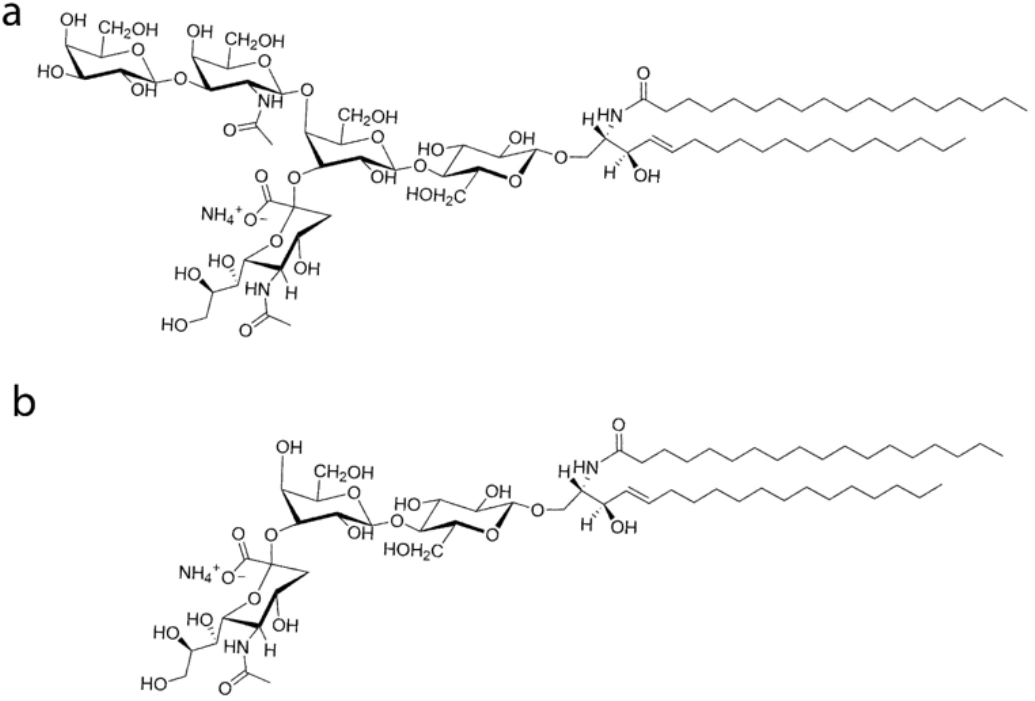
Representative structures of (a) GM1 and (b) GM3 gangliosides

**Figure S3:**
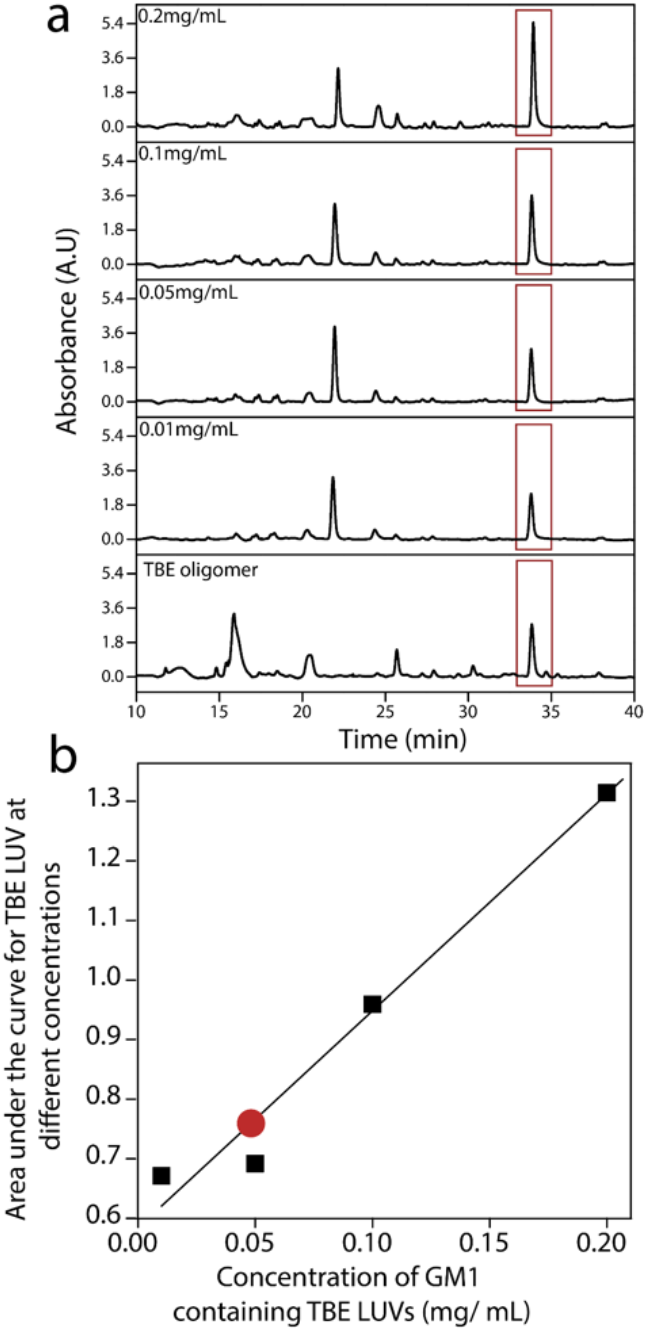
Quantification of lipid present within oligomers isolated from 50% GM1 enriched TBE LUV and Aβ reaction incubated for 5h. Standard curve for TBE lipid was plotted from area under the curve for HPLC chromatogram of TBE LUV at concentration of 0.2, 0.1, 0.05 and 0.01 mg/mL respectively. 5μM of isolated oligomer from 50% GM1 enriched TBE LUV and Aβ 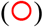 reaction was run on HPLC and peak corresponding to the TBE LUV was quantified using the standard curve plotted with TBE LUV.

**Table S1:**
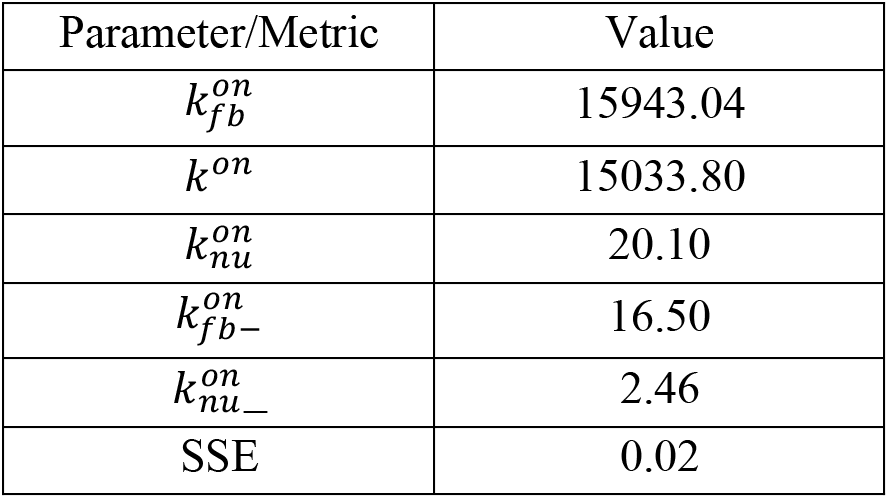
Table showing the parameter/metric values in case of the control data.

**Table S2:**
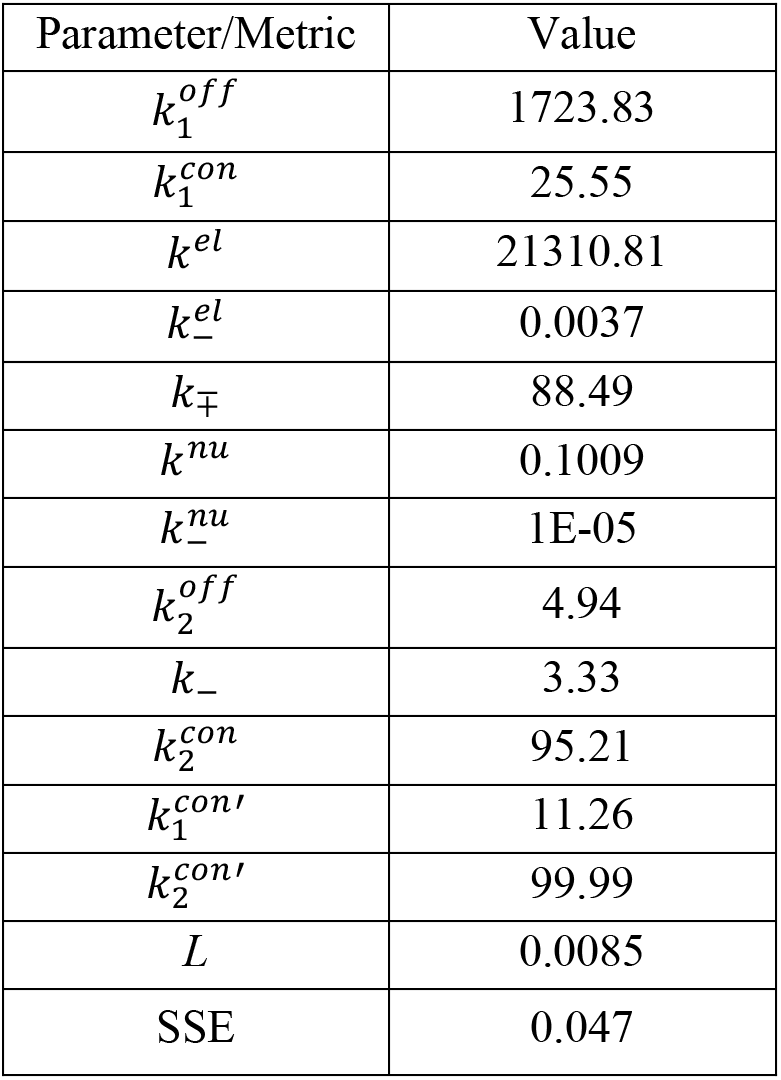
Table showing the parameter/metric values in case of the oligomerization data with 0% GM1 lipids and monomers.

**Table S3:**
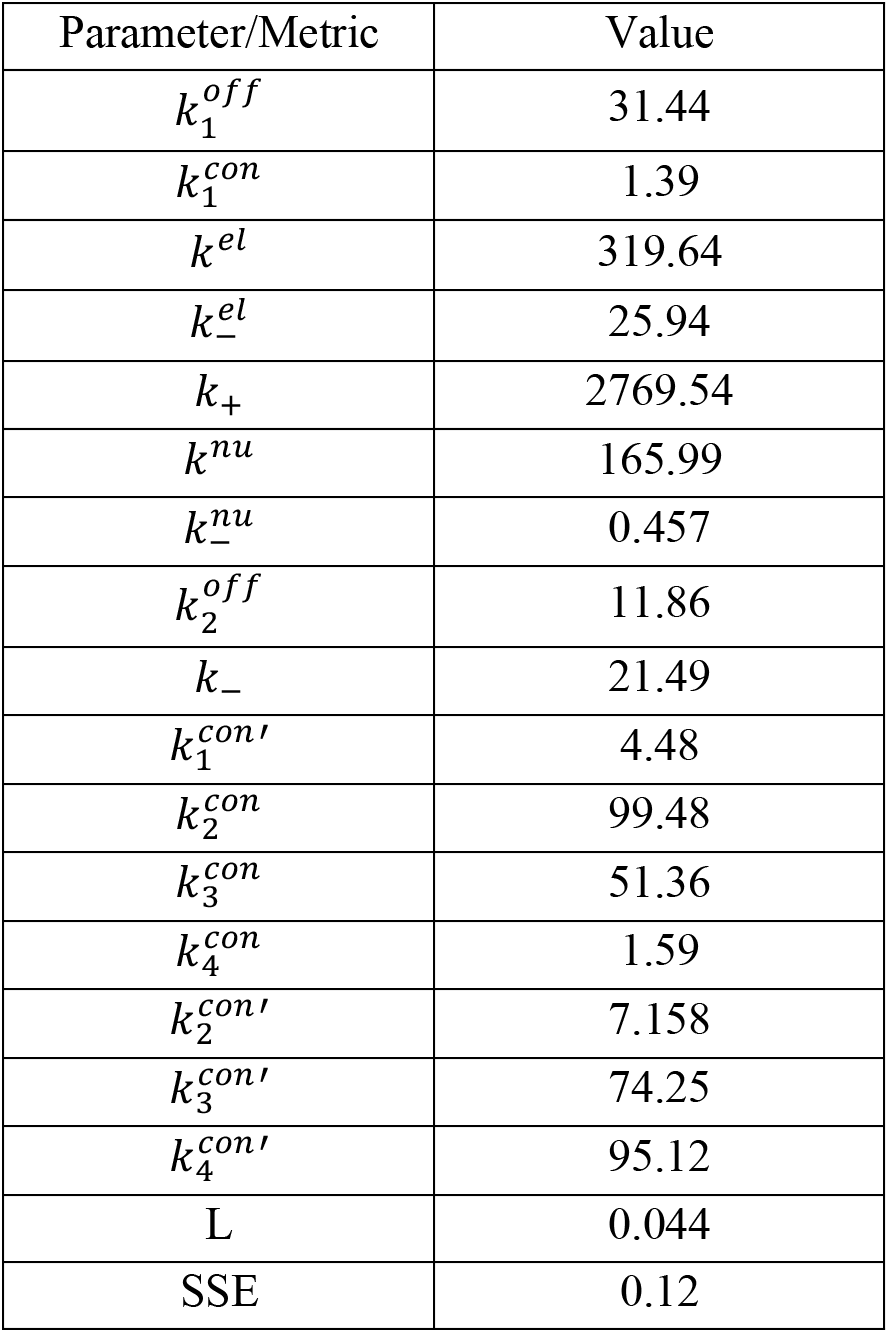
Table showing the parameter/metric values in case of the oligomerization data with 50GM1 lipids and monomers.

**Table S4:**
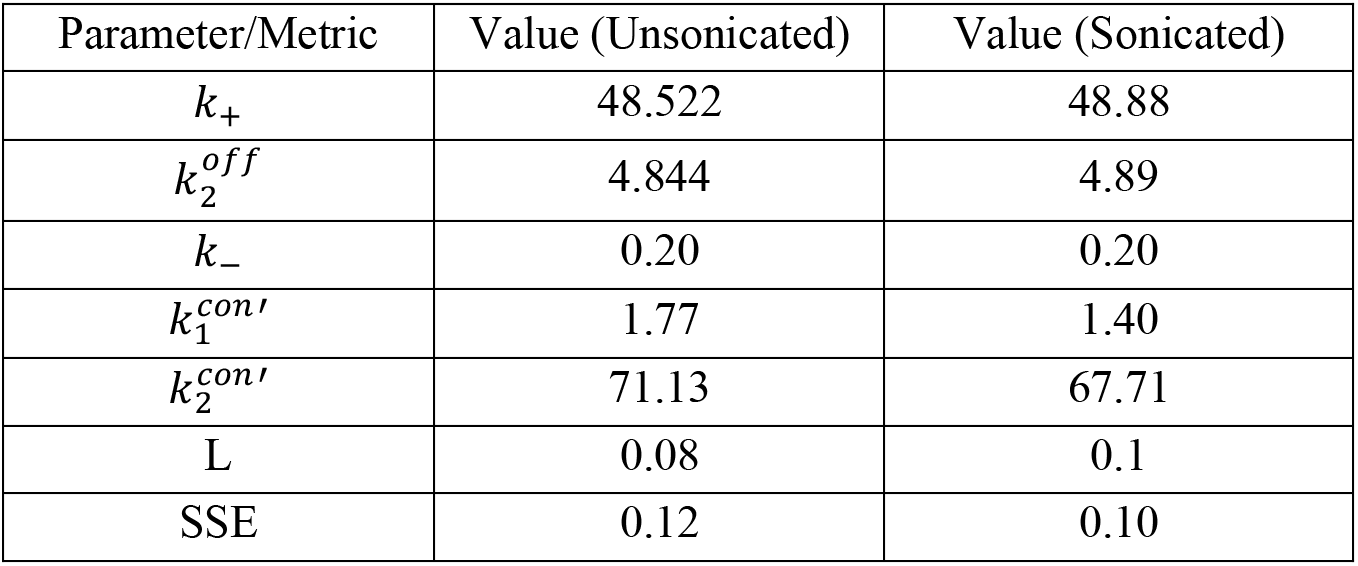
Table showing the parameter/metric values in case of the oligomerization data with 0% GM1 lipids and fibrils.

**Table S5:**
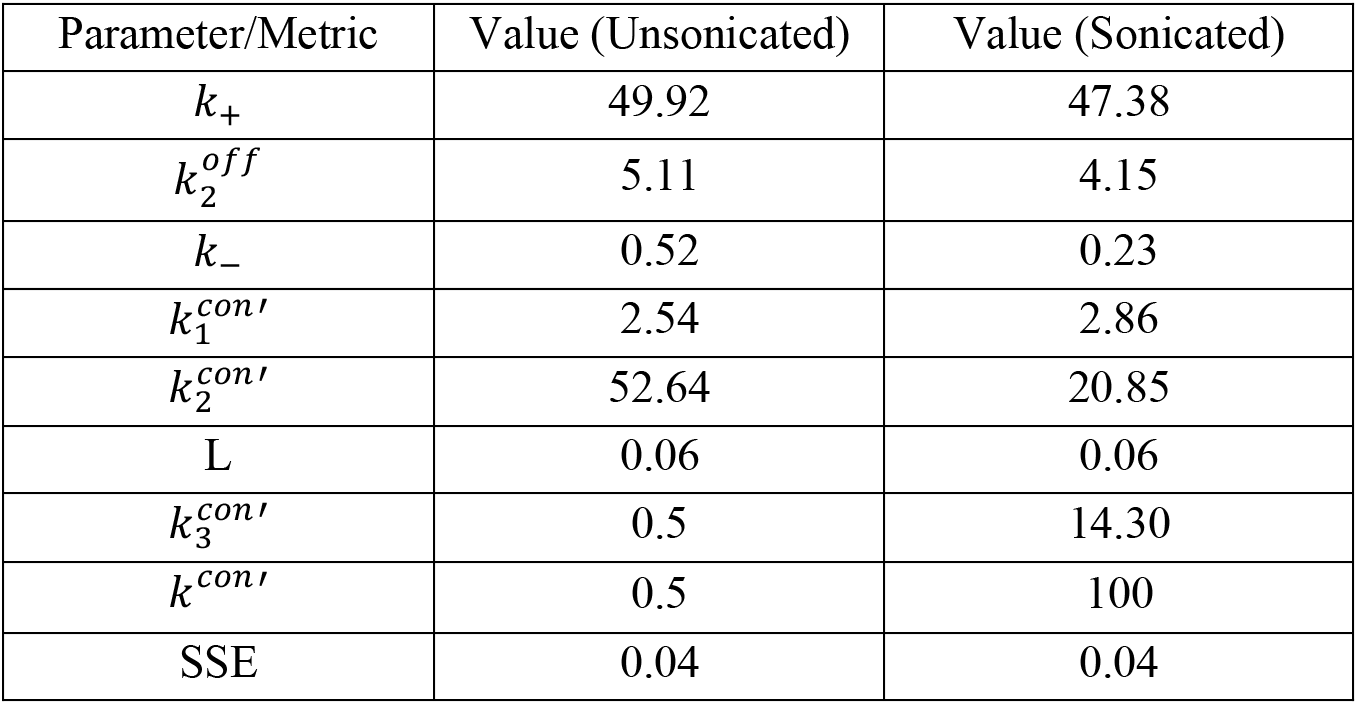
Table showing the parameter/metric values in case of the oligomerization data with 50% GM1 lipids and fibrils.

## Notes

### Competing Interest Statement

The authors have declared no competing interest.

### Summary of Updates

We have now corrected some figure numbering issues. We apologize for the oversight.

## REFERENCES

(1) Shoji, M., Golde, T. E., Ghiso, J., Cheung, T. T., Estus, S., Shaffer, L. M., Cai, X. D., McKay, D. M., Tintner, R., Frangione, B., and et, al. (1992) Production of the Alzheimer amyloid beta protein by normal proteolytic processing. Science (80-.). 258, 126 LP – 129.

(2) Selkoe, D. J., Podlisny, M. B., Joachim, C. L., Vickers, E. A., Lee, G., Fritz, L. C., and Oltersdorf, T. (1988) Beta-amyloid precursor protein of Alzheimer disease occurs as 110-to 135-kilodalton membrane-associated proteins in neural and nonneural tissues. Proc. Natl. Acad. Sci. 85, 7341–7345.

(3) Chow, V. W., Mattson, M. P., Wong, P. C., and Gleichmann, M. (2010) An overview of APP processing enzymes and products. Neuromolecular Med. 12, 1–12.

(4) Cohen, S. I. A., Linse, S., Luheshi, L. M., Hellstrand, E., White, D. A., Rajah, L., Otzen, D. E., Vendruscolo, M., Dobson, C. M., and Knowles, T. P. J. (2013) Proliferation of amyloid-β42 aggregates occurs through a secondary nucleation mechanism. Proc. Natl. Acad. Sci. 110, 9758 LP – 9763.

(5) Arosio, P., Knowles, T. P. J., and Linse, S. (2015) On the lag phase in amyloid fibril formation. Phys. Chem. Chem. Phys. 17, 7606–7618.

(6) Garai, K., and Frieden, C. (2013) Quantitative analysis of the time course of Aβ oligomerization and subsequent growth steps using tetramethylrhodamine-labeled Aβ. Proc. Natl. Acad. Sci. 110, 3321 LP – 3326.

(7) Li, S., and Selkoe, D. J. (2020) A mechanistic hypothesis for the impairment of synaptic plasticity by soluble Aβ oligomers from Alzheimer’s brain. J. Neurochem. 154, 583–597.

(8) Walsh, D. M., Klyubin, I., Fadeeva, J. V, Cullen, W. K., Anwyl, R., Wolfe, M. S., Rowan, M. J., and Selkoe, D. J. (2002) Naturally secreted oligomers of amyloid β protein potently inhibit hippocampal longterm potentiation in vivo. Nature 416, 535–539.

(9) Hong, W., Wang, Z., Liu, W., Malley, T. T. O., Jin, M., Willem, M., Haass, C., Frosch, M. P., and Walsh, D. M. (2018) Diffusible, highly bioactive oligomers represent a critical minority of soluble Aβ in Alzheimer ‘ s disease brain. Acta Neuropathol.

(10) Walsh, D. M., Townsend, M., Podlisny, M. B., Shankar, G. M., Fadeeva, J. V, Agnaf, O. El, Hartley, D. M., and Selkoe, D. J. (2005) Certain Inhibitors of Synthetic Amyloid β-Peptide (Aβ) Fibrillogenesis Block Oligomerization of Natural Aβ and Thereby Rescue Long-Term Potentiation. J. Neurosci. 25, 2455 LP – 2462.

(11) Stroud, J. C., Liu, C., Teng, P. K., and Eisenberg, D. (2012) Toxic fibrillar oligomers of amyloid-have cross-structure. Proc. Natl. Acad. Sci. 109, 7717–7722.

(12) De Felice, F. G., Velasco, P. T., Lambert, M. P., Viola, K., Fernandez, S. J., Ferreira, S. T., and Klein, W. L. (2007) Aβ oligomers induce neuronal oxidative stress through an N-methyl-D-aspartate receptor-dependent mechanism that is blocked by the Alzheimer drug memantine. J. Biol. Chem. 282, 11590–11601.

(13) Soto, C. (2003) Unfolding the role of protein misfolding in neurodegenerative diseases. Nat. Rev. Neurosci. 4, 49.

(14) Sciacca, M. F. M., Kotler, S. A., Brender, J. R., Chen, J., Lee, D., and Ramamoorthy, A. (2012) Two-Step Mechanism of Membrane Disruption by Aβ through Membrane Fragmentation and Pore Formation. Biophys. J. 103, 702–710.

(15) Rangachari, V., Dean, D. N., Rana, P., Vaidya, A., and Ghosh, P. (2018) Cause and consequence of Aβ — Lipid interactions in Alzheimer disease pathogenesis. Biochim. Biophys. Acta - Biomembr. 1860, 1652–1662.

(16) Rangachari, V., Moore, B. D., Reed, D. K., Sonoda, L. K., Bridges, A. W., Conboy, E., Hartigan, D., and Rosenberry, T. L. (2007) Amyloid-β(1-42) Rapidly Forms Protofibrils and Oligomers by Distinct Pathways in Low Concentrations of Sodium Dodecylsulfate. Biochemistry 46, 12451–12462.

(17) Kumar, A., Bullard, R. L., Patel, P., Paslay, L. C., Singh, D., Bienkiewicz, E. A., Morgan, S. E., and Rangachari, V. (2011) Non-Esterified Fatty Acids Generate Distinct Low-Molecular Weight Amyloid-β (Aβ42) Oligomers along Pathway Different from Fibril Formation. PLoS One (Ikezu, T., Ed.) 6, e18759.

(18) Dean, D. N., Das, P. K., Rana, P., Burg, F., Levites, Y., Morgan, S. E., Ghosh, P., and Rangachari, V. (2017) Strain-specific fibril propagation by an Aβ dodecamer. Sci. Rep. 7, 40787.

(19) Kumar, A., Paslay, L. C., Lyons, D., Morgan, S. E., Correia, J. J., and Rangachari, V. (2012) Specific soluble oligomers of amyloid-β peptide undergo replication, and form non-fibrillar aggregates In interfacial environments. J. Biol. Chem. jbc-M112.

(20) Khondker, A., Alsop, R. J., and Rheinstädter, M. C. (2017) Membrane-accelerated amyloid-β aggregation and formation of cross-β sheets. Membranes (Basel). 7, 49.

(21) Korshavn, K. J., Satriano, C., Lin, Y., Zhang, R., Dulchavsky, M., Bhunia, A., Ivanova, M. I., Lee, Y. H., La Rosa, C., Lim, M. H., and Ramamoorthy, A. (2017) Reduced lipid bilayer thickness regulates the aggregation and cytotoxicity of amyloid-β. J. Biol. Chem. 292, 4638–4650.

(22) Saha, J., Dean, D. N., Dhakal, S., Stockmal, K. A., Morgan, S. E., Dillon, K. D., Adamo, M. F., Levites, Y., and Rangachari, V. (2021) Biophysical characteristics of lipid-induced Aβ oligomers correlate to distinctive phenotypes in transgenic mice. FASEB J. 35.

(23) Terzi, E., Hölzemann, G., and Seelig, J. (1997) Interaction of Alzheimer β-amyloid peptide (1-40) with lipid membranes. Biochemistry 36, 14845–14852.

(24) Zhao, H., Tuominen, E. K. J., and Kinnunen, P. K. J. (2004) Formation of Amyloid Fibers Triggered by Phosphatidylserine-Containing Membranes ^†^. Biochemistry 43, 10302–10307.

(25) Ege, C., and Lee, K. Y. C. (2004) Insertion of Alzheimer’s Aβ40 peptide into lipid monolayers. Biophys. J. 87, 1732–1740.

(26) McLaurin, J., and Chakrabartty, A. (1997) Characterization of the interactions of Alzheimer β-amyloid peptides with phospholipid membranes. Eur. J. Biochem. 245, 355–363.

(27) Accardo, A., Shalabaeva, V., Cotte, M., Burghammer, M., Krahne, R., Riekel, C., and Dante, S. (2014) Amyloid β peptide conformational changes in the presence of a lipid membrane system. Langmuir 30, 3191–3198.

(28) Ke, P. C., Zhou, R., Serpell, L. C., Riek, R., Knowles, T. P. J., Lashuel, H. A., Gazit, E., Hamley, I. W., Davis, T. P., and Fändrich, M. (2020) Half a century of amyloids: past, present and future. Chem. Soc. Rev. 49, 5473–5509.

(29) Nicastro, M. C., Spigolon, D., Librizzi, F., Moran, O., Ortore, M. G., Bulone, D., Biagio, P. L. S., and Carrotta, R. (2016) Amyloid β-peptide insertion in liposomes containing GM1-cholesterol domains. Biophys. Chem. 208, 9–16.

(30) Yu, X., and Zheng, J. (2012) Cholesterol Promotes the Interaction of Alzheimer β-Amyloid Monomer with Lipid Bilayer. J. Mol. Biol. 421, 561–571.

(31) Hayashi, H., Kimura, N., Yamaguchi, H., Hasegawa, K., Yokoseki, T., Shibata, M., Yamamoto, N., Michikawa, M., Yoshikawa, Y., Terao, K., Matsuzaki, K., Lemere, C. A., Selkoe, D. J., Naiki, H., and Yanagisawa, K. (2004) A Seed for Alzheimer Amyloid in the Brain. J. Neurosci. 24, 4894 LP – 4902.

(32) Yanagisawa, K., Odaka, A., Suzuki, N., and Ihara, Y. (1995) GM1 ganglioside-bound amyloid β-protein (Aβ): A possible form of preamyloid in Alzheimer's disease. Nat. Med. 1, 1062.

(33) Matsuzaki, K., and Horikiri, C. (1999) Interactions of amyloid β-peptide (1-40) with ganglioside-containing membranes. Biochemistry 38, 4137–4142.

(34) Mori, K., Mahmood, M. I., Neya, S., Matsuzaki, K., and Hoshino, T. (2012) Formation of GM1 ganglioside clusters on the lipid membrane containing sphingomyeline and cholesterol. J. Phys. Chem. B 116, 5111–5121.

(35) Matsuzaki, K., Kato, K., and Yanagisawa, K. (2010) Aβ polymerization through interaction with membrane gangliosides. Biochim. Biophys. Acta - Mol. Cell Biol. Lipids 1801, 868–877.

(36) Flagmeier, P., De, S., Michaels, T. C. T., Yang, X., Dear, A. J., Emanuelsson, C., Vendruscolo, M., Linse, S., Klenerman, D., Knowles, T. P. J., and Dobson, C. M. (2020) Direct measurement of lipid membrane disruption connects kinetics and toxicity of Aβ42 aggregation. Nat. Struct. Mol. Biol. 27, 886–891.

(37) Matsuzaki, K. (2020) Aβ—ganglioside interactions in the pathogenesis of Alzheimer’s disease. Biochim. Biophys. Acta (BBA)-Biomembranes 1862, 183233.

(38) Lin, H., Zhu, Y. J., and Lal, R. (1999) Amyloid β Protein (1-40) Forms Calcium-Permeable, Zn2+-Sensitive Channel in Reconstituted Lipid Vesicles,. Biochemistry 38, 11189–11196.

(39) Korshavn, K. J., Bhunia, A., Lim, M. H., and Ramamoorthy, A. (2016) Amyloid-β adopts a conserved, partially folded structure upon binding to zwitterionic lipid bilayers prior to amyloid formation. Chem. Commun. 52, 882–885.

(40) Zou, Y., Li, Y., Hao, W., Hu, X., and Ma, G. (2013) Parallel β-sheet fibril and antiparallel β-sheet oligomer: New insights into amyloid formation of hen egg white lysozyme under heat and acidic condition from FTIR spectroscopy. J. Phys. Chem. B 117, 4003–4013.

(41) Byler, D. M., and Susi, H. (1986) Examination of the secondary structure of proteins by deconvolved FTIR spectra. Biopolym. Orig. Res. Biomol. 25, 469–487.

(42) Susi, H., and Byler, D. M. (1986) [13] Resolution-enhanced fourier transform infrared spectroscopy of enzymes, in Methods in enzymology, pp 290–311. Elsevier.

(43) Das, P. K., Dean, D. N., Fogel, A. L., Liu, F., Abel, B. A., McCormick, C. L., Kharlampieva, E., Rangachari, V., and Morgan, S. E. (2017) Aqueous RAFT synthesis of glycopolymers for determination of saccharide structure and concentration effects on amyloid β aggregation. Biomacromolecules 18, 3359–3366.

(44) Bristol, A. N., Saha, J., George, H. E., Das, P. K., Kemp, L. K., Jarrett, W. L., Rangachari, V., and Morgan, S. E. (2020) Effects of Stereochemistry and Hydrogen Bonding on Glycopolymer-Amyloid-β Interactions. Biomacromolecules 21.

(45) Glover, F. (1986) Future paths for integer programming and links to artificial intelligence. Comput. Oper. Res. 13, 533–549.

(46) Rana, P., Bose, P., Vaidya, A., Rangachari, V., and Ghosh, P. (2020) Global fitting and parameter identifiability for amyloid-β aggregation with competing pathways, in 2020 IEEE 20th International Conference on Bioinformatics and Bioengineering (BIBE), pp 73–78.

(47) Condello, C., Lemmin, T., Stöhr, J., Nick, M., Wu, Y., Maxwell, A. M., Watts, J. C., Caro, C. D., Oehler, A., Keene, C. D., Bird, T. D., Duinen, S. G. van, Lannfelt, L., Ingelsson, M., Graff, C., Giles, K., DeGrado, W. F., and Prusiner, S. B. (2018) Structural heterogeneity and intersubject variability of Aβ in familial and sporadic Alzheimer’s disease. Proc. Natl. Acad. Sci. 115, E782–E791.

(48) Qiang, W., Yau, W.-M., Lu, J.-X., Collinge, J., and Tycko, R. (2017) Structural variation in amyloid-β fibrils from Alzheimer’s disease clinical subtypes. Nature 541, 217.

(49) Petkova, A. T., Leapman, R. D., Guo, Z., Yau, W.-M., Mattson, M. P., and Tycko, R. (2005) Self-propagating, molecular-level polymorphism in Alzheimer’s ß-amyloid fibrils. Science (80-.). 307, 262–265.

(50) Huang, D., Zimmerman, M. I., Martin, P. K., Nix, A. J., Rosenberry, T. L., and Paravastu, A. K. (2015) Antiparallel β-sheet structure within the C-terminal region of 42-residue Alzheimer’s amyloid-β peptides when they form 150-kDa oligomers. J. Mol. Biol. 427, 2319–2328.

(51) Lu, J.-X., Qiang, W., Yau, W.-M., Schwieters, C. D., Meredith, S. C., and Tycko, R. (2013) Molecular structure of β-amyloid fibrils in Alzheimer’s disease brain tissue. Cell 154, 1257–1268.

(52) Paravastu, A. K., Leapman, R. D., Yau, W.-M., and Tycko, R. (2008) Molecular structural basis for polymorphism in Alzheimer’s β-amyloid fibrils. Proc. Natl. Acad. Sci. 105, 18349–18354.

(53) Paravastu, A. K., Qahwash, I., Leapman, R. D., Meredith, S. C., and Tycko, R. (2009) Seeded growth of β-amyloid fibrils from Alzheimer’s brain-derived fibrils produces a distinct fibril structure. Proc. Natl. Acad. Sci. 106, 7443–7448.

(54) Dean, D. N., Rana, P., Campbell, R. P., Ghosh, P., and Rangachari, V. (2018) Propagation of an Aβ Dodecamer Strain Involves a Three-Step Mechanism and a Key Intermediate. Biophys. J. 114, 539–549.

(55) Yip, C. M., and McLaurin, J. (2001) Amyloid-β Peptide Assembly: A Critical Step in Fibrillogenesis and Membrane Disruption. Biophys. J. 80, 1359–1371.

(56) Lau, T.-L., Ambroggio, E. E., Tew, D. J., Cappai, R., Masters, C. L., Fidelio, G. D., Barnham, K. J., and Separovic, F. (2006) Amyloid-β Peptide Disruption of Lipid Membranes and the Effect of Metal Ions. J. Mol. Biol. 356, 759–770.

(57) Kotler, S. A., Walsh, P., Brender, J. R., and Ramamoorthy, A. (2014) Differences between amyloid-β aggregation in solution and on the membrane: insights into elucidation of the mechanistic details of Alzheimer’s disease. Chem. Soc. Rev. 43, 6692–6700.

(58) Vander Zanden, C. M., Wampler, L., Bowers, I., Watkins, E. B., Majewski, J., and Chi, E. Y. (2019) Fibrillar and nonfibrillar amyloid beta structures drive two modes of membrane-mediated toxicity. Langmuir 35, 16024–16036.

(59) Delgado, D. A., Doherty, K., Cheng, Q., Kim, H., Xu, D., Dong, H., Grewer, C., and Qiang, W. (2016) Distinct membrane disruption pathways are induced by 40-residueβ-Amyloid peptides. J. Biol. Chem. 291, 12233–12244.

(60) Qiang, W., Yau, W.-M., and Schulte, J. (2015) Fibrillation of β amyloid peptides in the presence of phospholipid bilayers and the consequent membrane disruption. Biochim. Biophys. Acta (BBA)-Biomembranes 1848, 266–276.

(61) Sabaté, R., Espargaró, A., Barbosa-Barros, L., Ventura, S., and Estelrich, J. (2012) Effect of the surface charge of artificial model membranes on the aggregation of amyloid β-peptide. Biochimie 94, 1730–1738.

(62) Sugiura, Y., Ikeda, K., and Nakano, M. (2015) High Membrane Curvature Enhances Binding, Conformational Changes, and Fibrillation of Amyloid-β on Lipid Bilayer Surfaces. Langmuir 31, 11549–11557.

(63) Terakawa, M. S., Lin, Y., Kinoshita, M., Kanemura, S., Itoh, D., Sugiki, T., Okumura, M., Ramamoorthy, A., and Lee, Y.-H. (2018) Impact of membrane curvature on amyloid aggregation. Biochim. Biophys. Acta (BBA)-Biomembranes.

(64) Lee, J., Kim, Y. H., T. Arce, F., Gillman, A. L., Jang, H., Kagan, B. L., Nussinov, R., Yang, J., and Lal, R. (2017) Amyloid β Ion Channels in a Membrane Comprising Brain Total Lipid Extracts. ACS Chem. Neurosci. 8, 1348–1357.

(65) Jang, H., Arce, F. T., Ramachandran, S., Capone, R., Lal, R., and Nussinov, R. (2010) β-Barrel Topology of Alzheimer’s β-Amyloid Ion Channels. J. Mol. Biol. 404, 917–934.

(66) Qi, J.-S., and Qiao, J.-T. (2001) Amyloid β-protein fragment 31-35 forms ion channels in membrane patches excised from rat hippocampal neurons. Neuroscience 105, 845–852.

(67) Chi, E. Y., Frey, S. L., and Lee, K. Y. C. (2007) Ganglioside GM1-Mediated Amyloid-beta Fibrillogenesis and Membrane Disruption. Biochemistry 46, 1913–1924.

(68) Okada, Y., Okubo, K., Ikeda, K., Yano, Y., Hoshino, M., Hayashi, Y., Kiso, Y., Itoh-Watanabe, H., Naito, A., and Matsuzaki, K. (2019) Toxic Amyloid Tape: A Novel Mixed Antiparallel/Parallel β-Sheet Structure Formed by Amyloid β-Protein on GM1 Clusters. ACS Chem. Neurosci. 10, 563–572.

(69) Wakabayashi, M., Okada, T., Kozutsumi, Y., and Matsuzaki, K. (2005) GM1 ganglioside-mediated accumulation of amyloid β-protein on cell membranes. Biochem. Biophys. Res. Commun. 328, 1019–1023.

(70) Kakio, A., Nishimoto, S., Yanagisawa, K., Kozutsumi, Y., and Matsuzaki, K. (2001) Cholesterol-dependent formation of GM1 ganglioside-bound amyloid β-protein, an endogenous seed for Alzheimer amyloid. J. Biol. Chem. 276, 24985–24990.

(71) Yoo, S., Zhang, S., Kreutzer, A. G., and Nowick, J. S. (2018) An Efficient Method for the Expression and Purification of Aβ(M1—42). Biochemistry 57, 3861–3866.

(72) Vivekanandan, S., Brender, J. R., Lee, S. Y., and Ramamoorthy, A. (2011) A partially folded structure of amyloid-beta(1-40) in an aqueous environment. Biochem. Biophys. Res. Commun. 411, 312–316.

(73) MacDonald, R. C., MacDonald, R. I., Menco, B. P. M., Takeshita, K., Subbarao, N. K., and Hu, L. (1991) Small-volume extrusion apparatus for preparation of large, unilamellar vesicles. Biochim. Biophys. Acta (BBA)-Biomembranes 1061, 297–303.

(74) Kremer, J. J., and Murphy, R. M. (2003) Kinetics of adsorption of β-amyloid peptide Aβ(1—40) to lipid bilayers. J. Biochem. Biophys. Methods 57, 159–169.

(75) Jimah, J. R., Schlesinger, P. H., and Tolia, N. H. (2017) Liposome Disruption Assay to Examine Lytic Properties of Biomolecules. Bio-protocol 7, e2433.

(76) Dhakal, S., Saha, J., Wyant, C. E., and Rangachari, V. (2021) αS Oligomers Generated from Interactions with a Polyunsaturated Fatty Acid and a Dopamine Metabolite Differentially Interact with Aβ to Enhance Neurotoxicity. ACS Chem. Neurosci. 12, 4153–4161.

(77) Saha, A., Mondal, G., Biswas, A., Chakraborty, I., Jana, B., and Ghosh, S. (2013) In vitro reconstitution of a cell-like environment using liposomes for amyloid beta peptide aggregation and its propagation. Chem. Commun. 49, 6119–6121.

(78) Rana, P., Dean, D. N., Steen, E. D., Vaidya, A., Rangachari, V., and Ghosh, P. (2017) Fatty Acid Concentration and Phase Transitions Modulate Aβ Aggregation Pathways. Sci. Rep. 7, 10370.

(79) Ghosh, P., Vaidya, A., Kumar, A., and Rangachari, V. (2016) Determination of critical nucleation number for a single nucleation amyloid-β aggregation model. Math. Biosci. 273, 70–79.

