## Supplementary material for "Ganglioside enriched phospholipid vesicles induce cooperative Aβ oligomerization and membrane disruption": none

Running title: *Aβ oligomerization in presence of liposomal membrane model system*

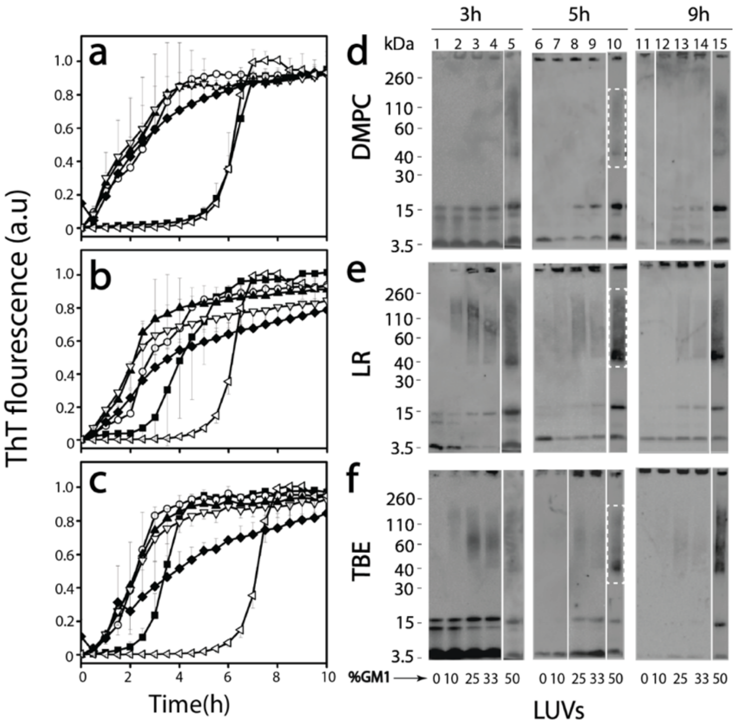


**Figure 1:** (a, b, and c) Normalized ThT fluorescence kinetics of buffered 25 μM Aβ without (◁; control) or with DMPC (a), LR (b) and TBE (c) LUVs each of them enriched with 10 (○), 25 (▲), 33 (▽), and 50 (◆) % GM1 ganglioside (by wt.) or without (■) GM1 in presence of 50 mM NaCl in 20mM tris buffer pH 8.00. (d, e, and f) Partially denaturing SDS-PAGE immunoblots of 25 μM Aβ in presence of DMPC, LR and TBE LUVs respectively, enriched without or with 10, 25, 33, and 50% GM1. Gels were run at intervals of 3, 5 and 9 hours, respectively.

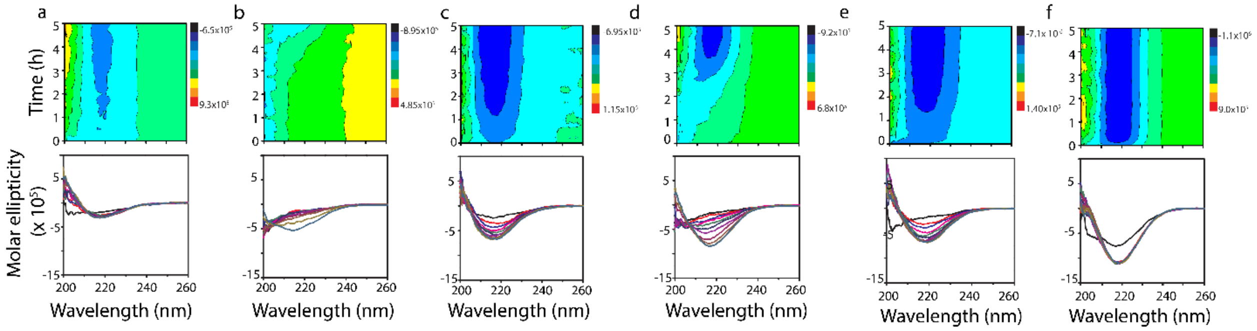


**Figure 2.** Far-UV CD contour and time course plots for buffered (20 mM tris buffer pH 8.0, 50 mM NaCl) 25 μM monomeric Aβ incubated with LUVs of 50 and 0% GM1 enriched DMPC (a and b, respectively), 50 and 0% GM1 enriched LR (c and d, respectively), and 50 and 0% GM1 enriched TBE (e and f, respectively) collected for up to five hours at 37 °C in quiescent conditions.

***Aβ oligomers isolated from GM1 enriched vesicles show distinctive biophysical characteristics.***

To see if oligomers generated in the presence are isolable, freshly purified Aβ monomers (25 μM) were incubated with 50% GM1 LUVs (DMPC, LR and TBE respectively) at 37 °C quiescent conditions. To isolate the oligomeric species from the reactions containing monomeric or fibrillar species, samples after 5 hours were centrifuged at 18000xg for 20 minutes. The supernatant was then subjected to fractionation by size exclusion chromatography (SEC) using a Superdex-75 column. Fractionation of all three LUVs showed two peaks near the void volume with a small first peak at fraction 15 and a larger one between fractions 16 and 18 (solid line; Figure 3c). In addition, a third peak at an included volume at fraction 24 was observed (solid line; Figure 3c).

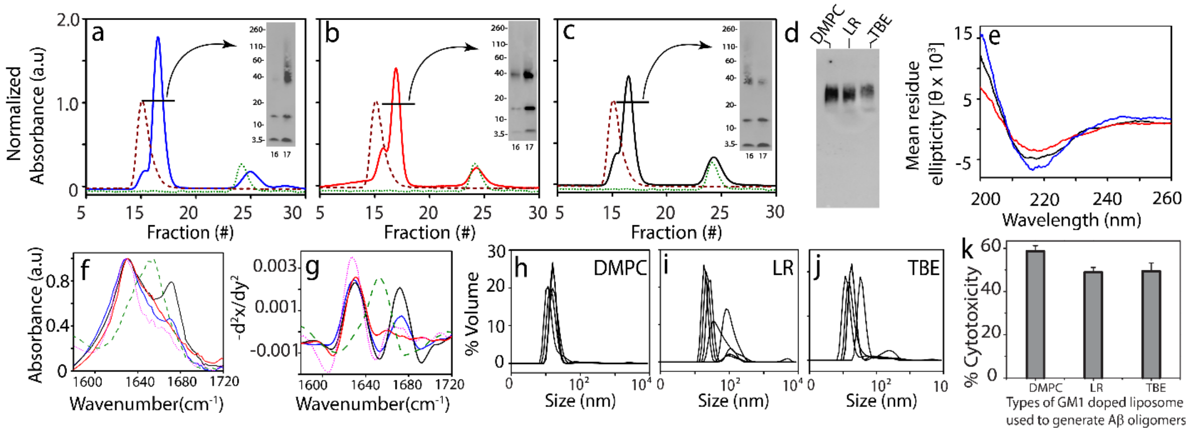

**Figure 3** (a-c) SEC chromatogram for isolation of Aβ oligomers generated in presence of 50% GM1 enriched DMPC (—), LR (—)and TBE LUVs (—) respectively , LUV control at 0.3 mg/mL (—) and control Aβ (—) at 5 h, inset- SDS PAGE immunoblots of SEC isolated oligomer fraction 16-17 (d) Native PAGE immunoblot for SEC isolated Aβ oligomers generated in the presence of 50% GM1 enriched DMPC, LR and TBE LUVs respectively, (e) CD spectra of fraction 17 of SEC isolated Aβ oligomers generated in the presence of 50% GM1 enriched DMPC (—), LR(—) and TBE LUVs(—) respectively (f) FTIR spectra for SEC isolated Aβ oligomers generated in the presence of 50% GM1 enriched TBE (—), DMPC (—) and LR (—) LUVs, homotypic Aβ fibril (—) and BSA control (—) respectively (g) Negative of double derivative of the FTIR spectra (fig3. (f)) (h-j) DLS for fraction 17 of SEC isolated Aβ oligomers generated in the presence of 50% GM1 enriched DMPC, LR and TBE LUVs respectively, (k) XTT assay performed on SHY5Y neuroblastoma cells upon incubation with isolated Aβ oligomers from 50% GM1 enriched DMPC, LR and TBE LUVs respectively expressed in terms of % of dead cells. n=3 independent cell cultures on isolated oligomers, statistically significant at $p < 0.05$ based on one-way ANOVA analysis.

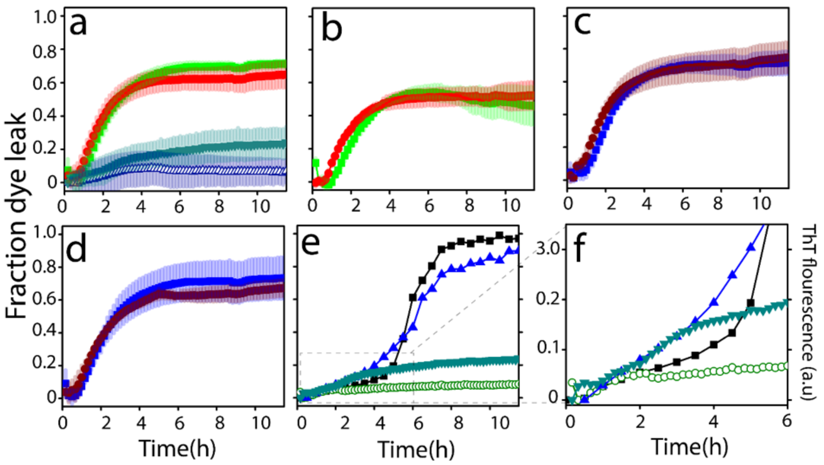


**Figure 4**: Vesicle dye leak analysis monitored by 6-carboxyflourescein (6-FAM) dye on; (a) TBE LUVs incubated with 10 μM Aβ monomers (△) or 2 μM isolated Aβ fibrils generated from the same liposomes (■); 50% GM1-enriched TBE LUVs incubated with 10μM Aβ monomers (▼); or 2 μM isolated Aβ fibrils generated from 50% GM1-enriched liposomes (●); (b) TBE LUVs incubated with 2 μM sonicated Aβ fibrils generated in the presence TBE liposomes (■) or 50% GM1-enriched TBE LUVs incubated with 2 μM sonicated Aβ fibrils generated in the presence of 50% GM1-enriched liposomes (●); (c) TBE LUVs incubated with 2 μM isolated Aβ fibrils generated in the absence of liposomes (■) or 50% GM1-enriched LUVs incubated with 2 μM isolated Aβ fibrils generated in the absence of liposomes (●); (d) samples in (c) but sonicated; (e) ThT fluorescence of 10μM Aβ monomers in presence of 50% GM1-enriched TBE LUVs (◀) and 50% GM3-enriched TBE LUVs (■); 6-FAM dye leakage of 50% GM1- enriched TBE LUVs (▼) and 50% GM3-enriched TBE LUVs (○) in presence of 10μM Aβ monomers; (f) zoomed-in image of figure 4(e) showing the initial 6 h of the reaction.

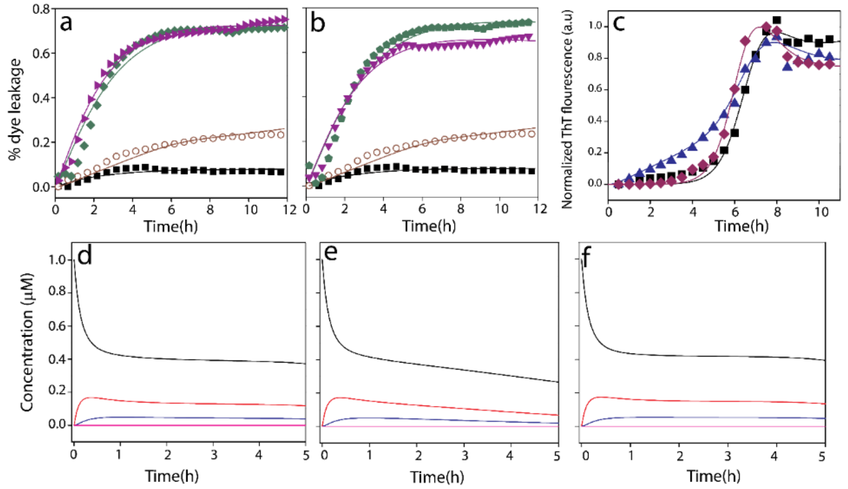


**Figure 5**: Computational fits of 6-carboxyflourescein dye leak assay of TBE LUVs (a) with 50% GM1 and 10 μM Aβ monomers (○), without GM1 and 10 μM Aβ monomers (■), with 50% GM1 and 2 μM Aβ fibrils (▶), without GM1 and 2 μM Aβ fibril (◆) (b) with 50% GM1 and 10 μM Aβ monomers (○), without GM1 and 10 μM Aβ monomers (■), with 50% GM1 and 2 μM sonicated Aβ fibrils (▼), without GM1 and 2 μM sonicated Aβ fibril (⬟) (c) Normalized ThT fluorescence kinetics of buffered 10 μM Aβ without (■; control) or with TBE LUVs each of them enriched with 50 (▲) % GM1 ganglioside (by wt.) or without (◆) GM1 in presence of 50 mM NaCl in 10mM sodium phosphate buffer pH 8.00; Aβ monomers (A1(−)), oligomers (A2 (−) and A3 (−)) and fibrils (F (−)) distribution plots for first 5 h from the start of reactions of Aβ monomers with TBE LUVs, (d) no GM1, (e) 50% GM1 or (f) without LUVs.

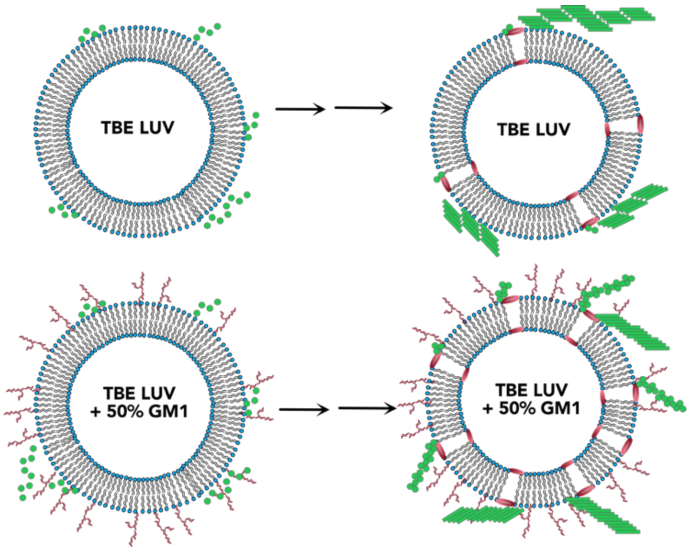


**Figure 6.** Schematic of conclusions drawn from this study showing the effect of GM1 enrichment in liposomes.

The study presented here shows alteration of surface characteristics, especially to the degree of charge density by dilution with neutral GM1 gangliosides seem to decisively affect the oligomerization of Aβ (Figure 6). While LUVs without or very low amount of GM1 accelerates the aggregation of Aβ to form higher molecular weight fibrils in the first five hours of incubation, LUVs enriched with high concentration of GM1 causes oligomerization of Aβ on the LUV surface, kinetically trapping the Aβ oligomers. Among the three different types of LUVs used, i.e., DMPC, LR and TBE that have different compositions, were found to augment aggregation of Aβ but also showed oligomerization when enriched with 50% GM1. This implicates the significance of gangliosides in Aβ aggregation as previous studies have established[35,67–70]. Furthermore, the GM1 enriched TBE LUVs showed somewhat modified ThT aggregation kinetics that correlated with a partially helical conformational state at an early aggregation stage. More importantly, these temporal changes also coincided with membrane disruption brought upon only by high GM1-enriched samples. This phenomenon may be due to altered lipid-packaging or dilution of anionic charge density or both due to GM1 enrichment. It is noteworthy that the pore formation was not abrupt but rather was slow and progressive in nature but only showed ~ 20% at the end of 11 hours of incubation (Figure 4). By contrast, fibrils and sonicated fibrils of Aβ generated in the presence and absence of liposome showed rapid pore formation. Furthermore, addition of fibrils generated from GM1-enriched liposomes too showed rapid pore formation. Two possible explanations can be rendered for these observations; *a*) the oligomers formed during aggregation is low in concentration as computed (~ 0.5 μM; Figure 5e) to effect rapid change in pore formation kinetics, or *b*) not oligomers but high molecular weight fibrils effect membrane disruption more effectively. A third explanation could be that the mechanisms of pore formation could be either numerous small pores or a few large pores for monomers aggregating on the surface or when pre-formed aggregates are added, respectively. Yet another key observation is that the oligomerization and membrane disruption is also selective to the nature of sugar distributions on gangliosides. While GM1 ganglioside promotes membrane pore formation, GM3 does seem to have such an effect, nor does it promote oligomers. Collectively, the data indicate that for early stages of oligomer formation, membrane selectivity is important, to generate conformationally distinct and toxic species, however at later stages when the higher molecular weight species are already formed, membrane is ruptured in a different mechanism than while Aβ oligomerization. Furthermore, oligomerization and pore formation seem to be cooperative and coupled to one another. As mentioned earlier, our lab and others have reported the formation of structurally distinct Aβ aggregates with equally distinct biophysical properties in presence of GM1 gangliosides. Recently, Matsuzaki and his group reported the formation of amyloid tape fibrils with mixed parallel and antiparallel β-sheet structure in presence of GM1 in membrane model systems[68]. Therefore, it can be concluded that membrane lipid composition as well as GM1 content plays a role in generating distinct oligomers. This inference is further supported by our CD time course data, where secondary structure of the intermediate species along the pathway of oligomerization are different for liposomes enriched with GM1 gangliosides. In this report, we further these findings to uncover that GM1 ganglioside enrichment of membrane vesicles not only promotes oligomerization but also induces membrane disruption in a cooperative manner. This suggests that aggregation and modulation of membrane dynamics can be coupled to each other, one that is facilitated as a function of specific membrane compositions. Such cooperative mechanisms may lead to the generation of conformational oligomers, which we believe are key templates for the formation of polymorphic fibrils observed in patient brains.

$$A_i + A_1 \leftrightarrows A_{i+1};\ \forall\ i \in \{1,2,...,11\}\ \ (k_{nu}^{on}/k_{nu-}^{on}\{\textit{pre-nucleation reactions}\}$$

$$F + A_i \leftrightarrows F;\ \forall\ i \in \{1,2,...,11\}\ \ (k_{fb}^{on}/k_{fb-}^{on}\{\textit{post-nucleation reactions}\} \qquad (1)$$

To reduce the number of species considered in the on-pathway reactions, we abstracted all post-nucleation species ($A_{12}$ onwards) as on-pathway fibrils denoted by *F*. Here, the forward and backward rate constants ($k_{nu}^{on}$ and $k_{nu-}^{on}$, respectively) for all pre-nucleation reactions were considered the same to reduce the number of parameters and based on our prior work[18,78] . Similarly, the forward and backward rate constants ($k_{fb}^{on}$ and $k_{fb-}^{on}$, respectively) for all post-nucleation reactions were also considered the same. The intensity of the ThT data was mapped to the sum of the concentrations of the on-pathway fibrils as follows:

$$Int\ ThT^{on} = k^{on} * F$$

where, $k^{on}$ is a scaling constant used for fitting ThT intensity ($ThT^{on}$) to the fibril concentration.

*b) Reactions involving liposomes*. Firstly, the forward and the backward pre-nucleation and post-nucleation rate constants computed from the controls were used for the on-pathway species for oligomerization reactions (shown below), which facilitated the reduction of the number of estimable parameters in this phase. To model the oligomerization reactions, both the ThT aggregation kinetics data and the FITC dye-leak data were considered. In our models, the initial concentrations of the liposomes were varied as their molar mass could not be precisely calculated. A sequential, multiple pore formation model was considered as opposed to one expanding pore although both are possible; since we do not have enough evidence to discount one over the other, we chose the former arbitrarily. Two possible scenarios were considered based on experimental evidence: pores formed by (*i*) a pre-nucleation oligomer ($A_j$) and/or (*ii*) on-pathway fibrils (i.e., a post-nucleated oligomer denoted as *F*). These were modeled for the first pore, $BA_i$ by;

$$A_j + L \leftrightarrows BA_j;\ (k_1^{con})$$

$$F + L \leftrightarrows BA_{12};\ (k_1^{con\prime})$$

while for subsequent holes ($CA_i$, $DA_i$, $EA_i$), this is modeled by reactions of type:

$$BA_{12} + A_j \leftrightarrows CA_j;\ (k_2^{con})$$

$$BA_{12} + F \leftrightarrows CA_{12};\ (k_2^{con\prime})$$

Here, $A_j$ denotes the minimum pre-nucleation oligomer that can form a pore and the value of the *i*mer was identified through our parameter fitting mechanism. Moreover, the values of the rate constant combination ($k_1^{con}$ *and* $k_1^{con\prime}$) can suggest which mechanism is more likely for the first pore formation (i.e., through pre-nucleation oligomer or post-nucleation fibrils); similarly, the rate constant combination ($k_2^{con}$ *and* $k_2^{con\prime}$) suggests which mechanism is more likely for the second pore formation and so on. The cooperativity between pore formation and aggregation was captured by considering further oligomerization reactions assisted by the edge of the pore up to 24mers denoted by reactions of type:

$$BA_i + A_1 \leftrightarrows BA_{i+1};\ \forall\ i \in \{j,...,23\}$$

We additionally consider a bulk oligomerization in the presence of fibrils by reactions of type:

$$BA_{12} + F \rightarrow BA_{24};$$

The cooperativity amongst pores is captured by ($k_1^{con}$ *and* $k_1^{con\prime}$) etc. For example, note that the second hole ($CA_i$) is formed only after the first pore is formed. This is ensured by reactions of type:

$$BA_{12} + A_j \leftrightarrows CA_j;\ (k_2^{con})$$

$$BA_{12} + F \rightarrow CA_{12};\ (k_2^{con\prime})$$

where, the presence of $BA_i$ is necessary for the formation of $C_i$. Additionally, if the rate constant pair ($k_1^{con}$ *and* $k_1^{con\prime}$)) is less than ($k_2^{con}$ *and* $k_2^{con\prime}$)), this will suggest higher cooperativity in hole formation; in other words, the second hole formation (controlled by $k_2^{con}$ *and* $k_2^{con\prime}$) is faster than the formation of the first hole. Finally, to map the concentration values to the ThT and FITC signals, we considered all the species weighted by the oligomer size for FITC signal (denoted by summations ranging from 1,…,24) while only weighted values of post-nucleated oligomers were considered in the ThT signal (denoted by summations ranging from 12,…,24).

The reactions considered for the oligomerization phase are as follows:

$$A_j + L \rightarrow B_j;\ (k_1^{con})\ \text{(j = 2 in case of 0\% GM1 liposomes and j = 3 in case of 50\% GM1 liposomes)}$$

$$B_i + A_1 \leftrightarrows BA_{i+1};\ \forall\ i \in \{j,...,23\}\ \ (k^{nu}/k_-^{nu}\ \text{till 12mer and}\ k^{el}/k_-^{el}\ \text{after that})$$

$$BA_{12} + F \leftrightarrows BA_{24};\ (k_+/k_-)$$

$$F + L \rightarrow BA_{12};\ (k_1^{con\prime}) \qquad (2)$$

where $BA_i$ denotes the first hole with an oligomer of size *i*-mers, and $L$ is the liposome.

$$BA_{12} + A_j \rightarrow C_j;\ (k_2^{con})\ \ \text{(j = 2 in case of 0\% GM1 and j = 3 in case of 50\% GM1)}$$

$$C_i + A_1 \leftrightarrows CA_{i+1};\ \forall\ i \in \{j,...,23\}\ \ (k^{nu}/k_-^{nu}\ \text{till 12mers and}\ k^{el}/k_-^{el}\ \text{after that})$$

$$CA_{12} + F \leftrightarrows CA_{24};\ (k_+/k_-)$$

$B_{12} + F \rightarrow C_{12}$; $(k_2^{con\prime})$ (3)

Fitting of data for 0% GM1 enriched liposomes were done only with the above reactions. Here, $C_i$ denotes the second hole with an $i$-mer. Similarly, for the following reactions, $D_i$ denotes the third hole, $E_i$ denotes the fourth hole and so on.

$C_{12} + A_j \rightarrow D_j$; $(k_3^{con})$ ) (j = 2 in case of 0% GM1 liposomes and j = 3 in case of 50% GM1 liposomes)

$D_i + A_1 \leftrightarrows D_{i+1}$; $\forall$ i $\in$ {j,...,23} $(k^{nu}/k_-^{nu}$ till 12mers and $k^{el}/k_-^{el}$ after that)

$D_{12} + F \leftrightarrows D_{24}$; $(k_+/k_-)$

$C_{12} + F \rightarrow D_{12}$; $(k_3^{con\prime}$ (4)

$D_{12} + A_j \rightarrow E_j$; $(k_4^{con})$ ) (j = 2 in case of 0% GM1 liposomes and j = 3 in case of 50% GM1 liposomes)

$E_i + A_1 \leftrightarrows E_{i+1}$; $\forall$ i $\in$ {j,...,23} $(k^{nu}/k_-^{nu}$ till 12mers and $k^{el}/k_-^{el}$ after that)

$E_{12} + F \leftrightarrows E_{24}$; $(k_+/k_-)$

$D_{12} + F \rightarrow E_{12}$; $(k_4^{con\prime}$ (5)

Fitting of data for 0% GM1 enriched liposomes were done only with the above reactions. i.e. four pores. Then ThT data were mapped to the sum of the concentrations of the on-pathway fibrils and all the off-pathway oligomers beyond 12mers as follows:

$Int\ ThT = Int\ ThT_{on} + k_1^{off} * (\sum_{i=12}^{24} i * BAi + \sum_{i=12}^{24} i * CAi)$, for the 0% GM1 enriched liposomes.

$Int\ ThT = Int\ ThT_{on} + k_1^{off} * (\sum_{i=12}^{24} i * BAi + \sum_{i=12}^{24} i * CAi + \sum_{i=12}^{24} i * DAi + \sum_{i=12}^{24} i * Ei)$, for the 50% GM1 enriched liposomes.

The FITC dye leak data were mapped to the concentration of the off-pathway oligomers as follows:

$Int\ FITC = k_2^{off} * (\sum_{i=2}^{24} i * BAi + \sum_{i=2}^{24} i * CAi)$, for 0% GM1 enriched liposomes.

$Int\ FITC = k_2^{off} * (\sum_{i=2}^{24} i * Bi * (\sum_{i=3}^{24} i * BAi + \sum_{i=3}^{24} i * CAi + \sum_{i=3}^{24} i * DAi + \sum_{i=3}^{24} i * EAi)$, for 50% GM1 enriched liposomes.

The model is extendable to any number of holes and the curve fitting with the experimental data infers the optimal number of holes to be considered. For example, for 0% GM1, we first experimented with four holes ($B_i$, $C_i$, $D_i$, $E_i$), which was then systematically reduced to one hole ($B_i$); in this case two holes gave the best global fit with ThT and FITC dye leak data.

**ACKNOWLEDGEMENTS**

The authors would like to thank the following agencies for their financial support: National Institute of Aging (1R56AG062292-01) and the National Science Foundation (NSF CBET 1802793) to VR. The authors also thank the National Center for Research Resources (5P20RR01647-11) and the National Institute of General Medical Sciences (8 P20 GM103476-11) from the National Institutes of Health for funding through INBRE for the use of their core facilities.

**AUTHOR INFORMATION**

### Corresponding Author

**Vijay Rangachari** – *Department of Chemistry and Biochemistry, School of Mathematics and Natural Sciences, and Center for Molecular and Cellular Biosciences, University of Southern Mississippi, Hattiesburg MS 39406, USA.*

### Authors

**Preetam Ghosh** – *Department of Computer Science, Virginia Commonwealth University, Richmond VA 23284*

**Jhinuk Saha** – *Department of Chemistry and Biochemistry, School of Mathematics and Natural Sciences, University of Southern Mississippi, Hattiesburg MS 39406, USA*

**Priyankar Bose** – *Department of Computer Science, Virginia Commonwealth University, Richmond VA 23284*

**Shailendra Dhakal** – *Department of Chemistry and Biochemistry, School of Mathematics and Natural Sciences, University of Southern Mississippi, Hattiesburg MS 39406, USA*

**TOC figure**

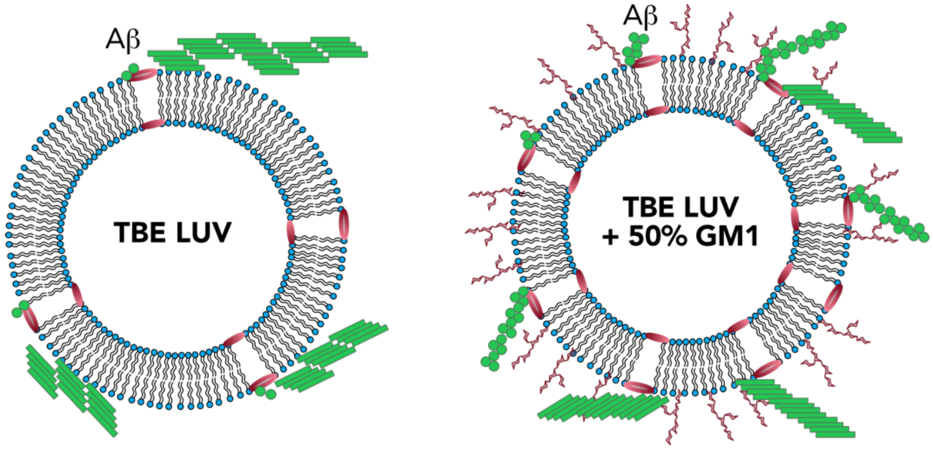


### SUPPLEMENTARY INFORMATION

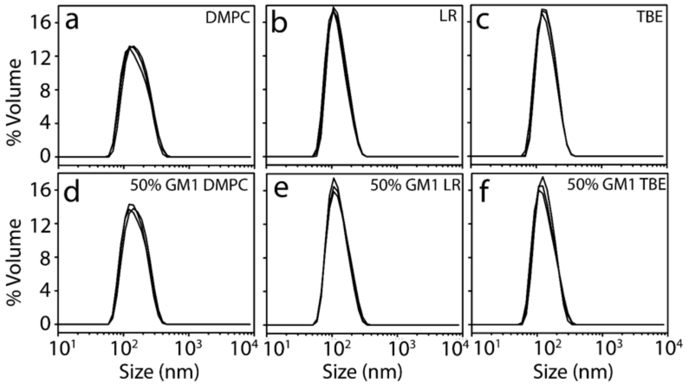


**Figure S1:** DLS of (0.2 mg/mL) DMPC, LR and TBE LUVs with (a,b, and c) or without (d,e, and f) GM1 extruded with 200 nm pore-sized polycarbonate membrane.

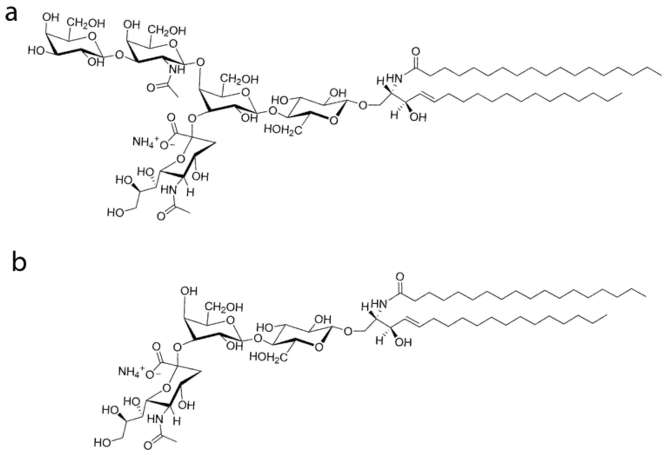


**Figure S2:** Representative structures of (a) GM1 and (b) GM3 gangliosides

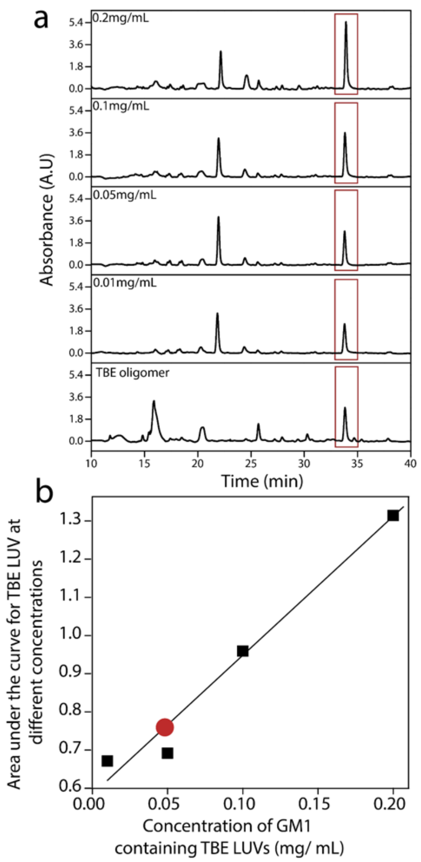


**Figure S3:** Quantification of lipid present within oligomers isolated from 50% GM1 enriched TBE LUV and Aβ reaction incubated for 5h. Standard curve for TBE lipid was plotted from area under the curve for HPLC chromatogram of TBE LUV at concentration of 0.2 ,0.1, 0.05 and 0.01 mg/mL respectively. 5µM of isolated oligomer from 50% GM1 enriched TBE LUV and Aβ (○) reaction was run on HPLC and peak corresponding to the TBE LUV was quantified using the standard curve plotted with TBE LUV.

| Parameter/Metric | Value |
|---|---|
| $k_{fb}^{on}$ | 15943.04 |
| $k^{on}$ | 15033.80 |
| $k_{nu}^{on}$ | 20.10 |
| $k_{fb-}^{on}$ | 16.50 |
| $k_{nu\_}^{on}$ | 2.46 |
| SSE | 0.02 |

**Table S1**: Table showing the parameter/metric values in case of the control data.

| Parameter/Metric | Value |
|---|---|
| $k_1^{off}$ | 1723.83 |
| $k_1^{con}$ | 25.55 |
| $k^{el}$ | 21310.81 |
| $k_-^{el}$ | 0.0037 |
| $k_\mp$ | 88.49 |
| $k^{nu}$ | 0.1009 |
| $k_-^{nu}$ | 1E-05 |
| $k_2^{off}$ | 4.94 |
| $k_-$ | 3.33 |
| $k_2^{con}$ | 95.21 |
| $k_1^{con\prime}$ | 11.26 |
| $k_2^{con\prime}$ | 99.99 |
| $L$ | 0.0085 |
| SSE | 0.047 |

**Table S2:** Table showing the parameter/metric values in case of the oligomerization data with 0% GM1 lipids and monomers.

| Parameter/Metric | Value |
|---|---|
| $k_1^{off}$ | 31.44 |
| $k_1^{con}$ | 1.39 |
| $k^{el}$ | 319.64 |
| $k_-^{el}$ | 25.94 |
| $k_+$ | 2769.54 |
| $k^{nu}$ | 165.99 |
| $k_-^{nu}$ | 0.457 |
| $k_2^{off}$ | 11.86 |
| $k_-$ | 21.49 |
| $k_1^{con\prime}$ | 4.48 |
| $k_2^{con}$ | 99.48 |
| $k_3^{con}$ | 51.36 |
| $k_4^{con}$ | 1.59 |
| $k_2^{con\prime}$ | 7.158 |
| $k_3^{con\prime}$ | 74.25 |
| $k_4^{con\prime}$ | 95.12 |
| L | 0.044 |
| SSE | 0.12 |

**Table S3:** Table showing the parameter/metric values in case of the oligomerization data with 50GM1 lipids and monomers.

| Parameter/Metric | Value (Unsonicated) | Value (Sonicated) |
|---|---|---|
| $k_+$ | 48.522 | 48.88 |
| $k_2^{off}$ | 4.844 | 4.89 |
| $k_-$ | 0.20 | 0.20 |
| $k_1^{con\prime}$ | 1.77 | 1.40 |
| $k_2^{con\prime}$ | 71.13 | 67.71 |
| L | 0.08 | 0.1 |
| SSE | 0.12 | 0.10 |

**Table S4:** Table showing the parameter/metric values in case of the oligomerization data with 0% GM1 lipids and fibrils.

| Parameter/Metric | Value (Unsonicated) | Value (Sonicated) |
|---|---|---|
| $k_+$ | 49.92 | 47.38 |
| $k_2^{off}$ | 5.11 | 4.15 |
| $k_-$ | 0.52 | 0.23 |
| $k_1^{con\prime}$ | 2.54 | 2.86 |
| $k_2^{con\prime}$ | 52.64 | 20.85 |
| L | 0.06 | 0.06 |
| $k_3^{con\prime}$ | 0.5 | 14.30 |
| $k^{con\prime}$ | 0.5 | 100 |
| SSE | 0.04 | 0.04 |

**Table S5:** Table showing the parameter/metric values in case of the oligomerization data with 50% GM1 lipids and fibrils.
